# How to deal with darkness: Modelling and visualization of zero-inflated personal light exposure data on a logarithmic scale

**DOI:** 10.1101/2024.12.30.630669

**Authors:** Johannes Zauner, Carolina Guidolin, Manuel Spitschan

## Abstract

Personal light exposure, the pattern of ocular light levels across time under free-living conditions measured with wearable devices, has become increasingly important in circadian and myopia research. Very small measurement values in light exposure patterns, especially zero, are regularly recorded in field studies. These zero-lux values are problematic for commonly applied logarithmic transformations, and should neither be dismissed nor be unduly influential in visualizations and statistical models. Common approaches used in zero-inflated data sets fail in at least one of these regards. We compare four ways to visualize such data on a linear, logarithmic, hybrid, or symlog scale and we model the light exposure patterns with a generalized additive model by removing zero-lux values, adding a very small or −1 log_10_ lux value to the dataset, or using the Tweedie error distribution. We show that a *symlog*-transformed visualization displays relevant features of light exposure across scales, including zero-lux, while at the same time reducing the emphasis on the small values (<1 lux). *Symlog* is well-suited to visualize differences in light exposure covering heavy-tailed negative values. The open-source software package *LightLogR* includes the symlog transformation for easy access. We further show that small but not negligible value additions to the light exposure data of -1 log_10_ lux for statistical modelling allow for acceptable models on a logarithmic scale, while very small values distort results. We also demonstrate the utility of the Tweedie distribution, which does not require prior transformations, models data on a logarithmic scale, and includes zero-lux values, capturing personal light exposure patterns satisfactorily. Data from field studies of personal light exposure requires appropriate handling of zero-lux values in a logarithmic context. *Symlog* scales for visualizations and an appropriate addition to input values for modelling, or the Tweedie distribution, provide a solid basis.

## Introduction

Personal light exposure, or the pattern of ocular light levels across time under free-living conditions, has become increasingly important in human health research [1]. In the past two decades, highly controlled laboratory research has shown the relevance of melanopsin-mediated pathways to the central pacemaker and various downstream effectors to regulate key physiological parameters important for well-being, alertness, sleep, and long-term mental and physical health [2, 3]. However, the real-world effects indicated by these insights can only be gauged when connecting the patterns of light people are exposed to under naturalistic conditions with relevant health outcomes. A growing literature of studies indicates that higher levels during the day and/or lower light levels at night are beneficial for sleep and mental and metabolic health [4-10], with more research required for different cultural, geographical, and other environmental conditions, as well as for different subgroups within a population [11, 12]. Environmental light levels are also relevant in the context of myopia development and progression [13-16]. Light exposure data in circadian and myopia research are commonly collected with small wearable devices attached to a person at eye level, to the chest, or, coming from actimetry research, placed on the wrist [17, 18].

Compared to controlled laboratory conditions, the dynamic range of light levels under free-living conditions ranges from 0 lux at night to more than 100,000 (10^5^) lux under bright daylight. This poses a unique measurement challenge for the light sensors in small wearable devices [17]. Particularly, this lower bound can be problematic from a measurement perspective and for the analysis, as complete darkness is not only possible but a regular occurrence in many people’s daily exposure patterns, e.g., such as in sleep environments. These circumstances make 0 lux a common value in datasets of personal light exposure, much more so than in past laboratory studies employing research-grade measurement equipment. The high dynamic range, in many cases, necessitates a logarithmic analysis, which is also appropriate since it mirrors how the retinal photoreceptors and downstream neural machinery operate [19, 20].

As zero cannot be logarithmically transformed into a real number, ‘zero’ instances require manual handling in both statistical models and visualizations. Currently, there are no standards for how these zero-lux values should be treated during data pre-processing. Without attention, these observations are often dropped – sometimes silently – from the analysis, skew results, and make plots ambiguous. We argue that treating small values outside the measurement range, particularly zero-lux values, should be intentional, deliberate, and transparent. This study takes a systematic approach to compare several strategies to treat zero lux illuminance values in personal light exposure datasets for statistical modelling and visualizations. We note that the relevance and methods shown here differ when using derived metrics based on illuminance measurements, i.e., not using the raw time series for visualization and modelling. While these metrics are highly relevant, analysis of the time series of measurements itself and the derived exposure patterns is becoming increasingly important, as it allows to model and explain individual environmental and behavioural influencing factors on personal light exposure [21]. Treating the zero-lux case deliberately in these scenarios is key.

## Method and Materials

The dataset used in this study is taken from Guidolin et al. (22), is available from Zenodo [23], and comes as part of the *LightLogR* software package [24]. It contains data for a week of personal light exposure measurement of a single participant and simultaneous environmental light measurements, taken from a university rooftop close to the participant at the horizontal plane and without any shading or obstructions. One exemplary day from this dataset is used for analysis. As the measure of personal light exposure, we use the melanopic equivalent of daylight illuminance (mel EDI) [25], which is part of the dataset. R statistical software [26] is used for analysis with the software package *LightLogR* (v0.4.2) [24] for data import and visualization. The visualization bases are taken from the *LightLogR* tutorial “The whole game” (https://tscnlab.github.io/LightLogR/articles/Day.html). The light exposure pattern is modelled with a generalized additive model [27] with the *mgcv* package (v1.9-1) [28], including the autoregressive error correction for lag 1 (AR1). The analysis documentation and the analysis source code are available as part of a Quarto HTML document as **Supplementary Material S1**.

### Zero-inflated data

We refer to zero-inflated data in the context of personal light exposure data. This term is most often used in the context of excess zero values for count data, where zeros exceed the prediction of a Poisson error model [29]. However, zero inflation is also appropriate in other contexts where observations of zero-lux are frequent and have to be treated with special care [29], which is the case in this topic.

### *Symlog* transformation

One of the scale transformations used for visualization is the *symlog* scale, or symmetrical log scale. *Symlog* is a logarithmic transformation allowing positive, negative, and zero input values. Positive values are logarithmically transformed. Negative values are logarithmically transformed through their absolute value, with a negative sign applied afterwards. A range between a freely defined positive value and its negative counterpart is excluded from logarithmic transformation. *Symlog* scales are available in Python as part of the *matplotlib* library [30] and in R as part of the *LightLogR* package [24].

### Smallest non-zero value

One of the treatments for non-zero values is to replace them with the smallest possible value so that 1+ x ≠ 1 (eq 1) is true. In R, this value is stored in the object *.Machine$double.eps* and is 2.20446 * 10^-16^ on the machine where the analysis was performed (MacBook Pro M4 Max, running macOS 15.1.1).

### Statistical modelling

**Table 1** summarizes the treatments of zero-lux measurements explored for statistical modelling.

**Table 1.** Model specifications. The modification of mel EDI specifies how input values to the model are transformed prior to modelling. The model formula specifies the input formula for the additive model in Wilkinson notation. type specifies that mel EDI can, on average, be different between the participant and the environment (daylight). s() denotes a specification of a smooth relationship between the dependent variable and the independent variable inside the brackets, time in this case, provided as seconds from midnight. The by = type specification fits a smooth relationship for each participant and environment. A basis spline bs = “cc” specifies a cyclic cubic regression spline, forcing the spline to connect on both ends in its value and its first and second derivative, thus taking into account the cyclic nature of daily light exposure. k = 24 specifies the knots, defining the maximum number of basis splines that add up to the smooth relationship.

| Model | Description | Modification of mel EDI | Model formula |
| --- | --- | --- | --- |
| 1 | Removing zero-lux observations | if mel EDI = 0<br>then mel EDI<sub>1</sub> = NA | $\log_{10}(\text{mel EDI}_1) \sim \text{type} + s(\text{time}, \text{by} = \text{type}, \text{bs} = "cc", k = 24)$ |
| 2 | Adding a very small, non-zero value | mel EDI<sub>2</sub> =<br>mel EDI<br>+ smallest float | $\log_{10}(\text{mel EDI}_2) \sim \text{type} + s(\text{time}, \text{by} = \text{type}, \text{bs} = "cc", k = 24)$ |
| 3 | Adding $-1 \log_{10}(\text{lux})$ | mel EDI<sub>3</sub> =<br>mel EDI + 0.1 | $\log_{10}(\text{mel EDI}_3) \sim \text{type} + s(\text{time}, \text{by} = \text{type}, \text{bs} = "cc", k = 24)$ |
| 4 | Using the Tweedie error distribution | mel EDI<sub>4</sub> = mel EDI | $\text{mel EDI}_4 \sim \text{type} + s(\text{time}, \text{by} = \text{type}, \text{bs} = "cc", k = 24)$ |

We chose these approaches with the following rationale:

- **Model 1:** This approach is most conservative from a measurement perspective, as these zero-lux values are almost certainly outside the measurement range of a wearable device.
- **Model 2:** This approach removes the issue of non-real logarithmically scaled values by adding the smallest possible value, thus changing the values minimally (∼2x10^-16^).
- **Model 3:** As Model 2, but without creating a gap in the value range on the logarithmic scale. 0.1 lux is -1 log_10_ lux below the lower measurement range specified by the manufacturer.
- **Model 4:** This approach does not include a treatment of the data but consists of the use of the Tweedie distribution [31], an error function that works with a logarithmic link function and allows zero and positive values.

## Results

### Visualizations

Figure 1 shows multiple ways to represent zero-lux measurements as part of a personal light exposure visualization. These are:

A. no scaling
B. logarithmic scaling
C. logarithmic scaling with a separate indicator for zero values (hybrid plot)
D. *symlog* scaling

**Figure 1.**
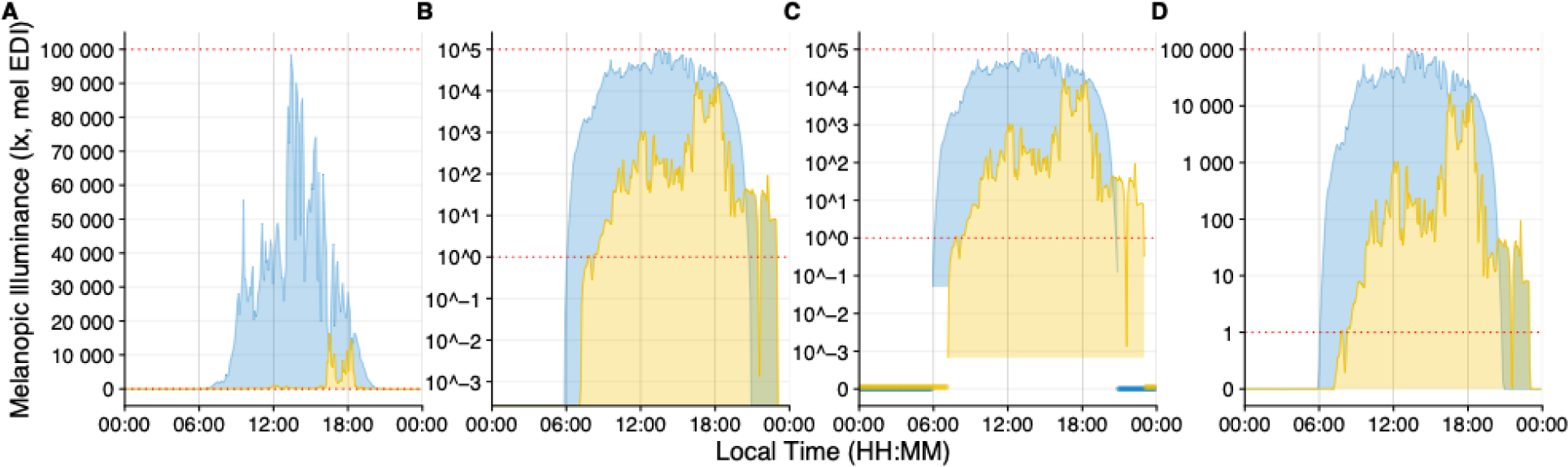
Scaling types to visualize light exposure patterns. All figures show melanopic EDI (lux) on the y-axis and local time on the x-axis. Yellow indicates the personal light exposure of a participant and blue the daylight potential in the area. Red dotted lines indicate the upper and lower measurement range provided by the manufacturer. **A**, Linear scaling. **B**, Logarithmic scaling with base 10. Zero-lux values are transformed to minus infinity and are not shown. **C**, Logarithmic scaling with base 10. Zero-lux values are shown at the bottom of the plot. The coloured ribbons of the plot indicate the minimum measurement value recorded by the device. **D**, *Symlog* scaling, with logarithmic scaling, base 10 above 1 lux, and linear scaling below.

Figure 2 shows multiple ways to visualize differences between different sources of light exposure. In this case this refers to the difference between environmental daylight conditions and a participants personal light exposure, however the principles are valid for differences of light exposure in general.

**Figure 2.**
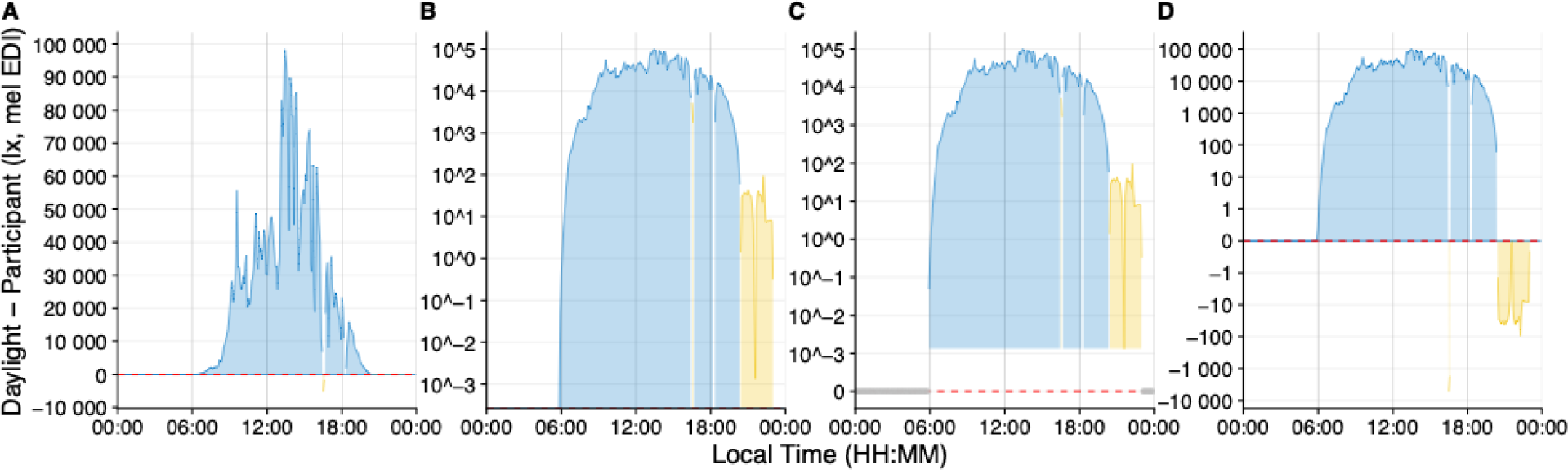
Scaling types to visualize differences in light exposure patterns. All figures show melanopic EDI (lux) on the y-axis and local time on the x-axis. Yellow indicates a participant’s personal light exposure value compared to daylight and blue vice versa. Red dashed lines indicate zero difference. **A**, Linear scaling. **B**, Logarithmic scaling with base 10. Zero-lux differences are transformed to minus infinity and are not shown. **C**, Logarithmic scaling with base 10. Zero-lux differences are shown at the bottom of the plot. The coloured ribbons of the plot indicate the minimum difference between the two patterns above zero. **D**, *Symlog* scaling, with logarithmic scaling, base 10 above 1 lux, no rescaling between -1 and 1, and base 10 below 1 (calculated from the absolute difference, followed by a sign flip).

### Statistical modelling

Figure 3 and Figure 4 show model results with the original data.

**Figure 3.**
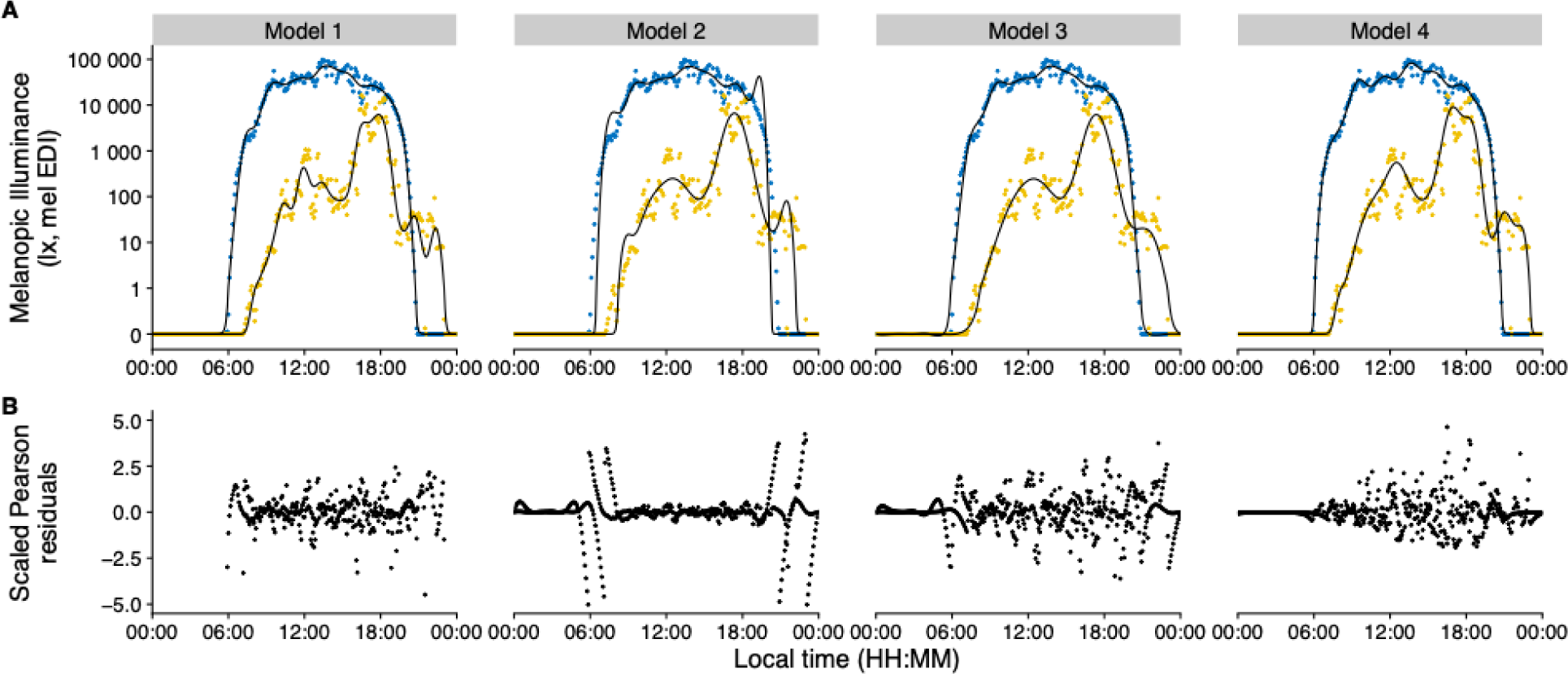
Model results and residual error distribution. **A,** Personal light exposure on a *symlog* scale for participant (yellow) and environmental light levels (blue). Points indicate measurements, black lines indicate the fitted model. **B,** Scaled Pearson residuals/errors for all models.

**Figure 4.**
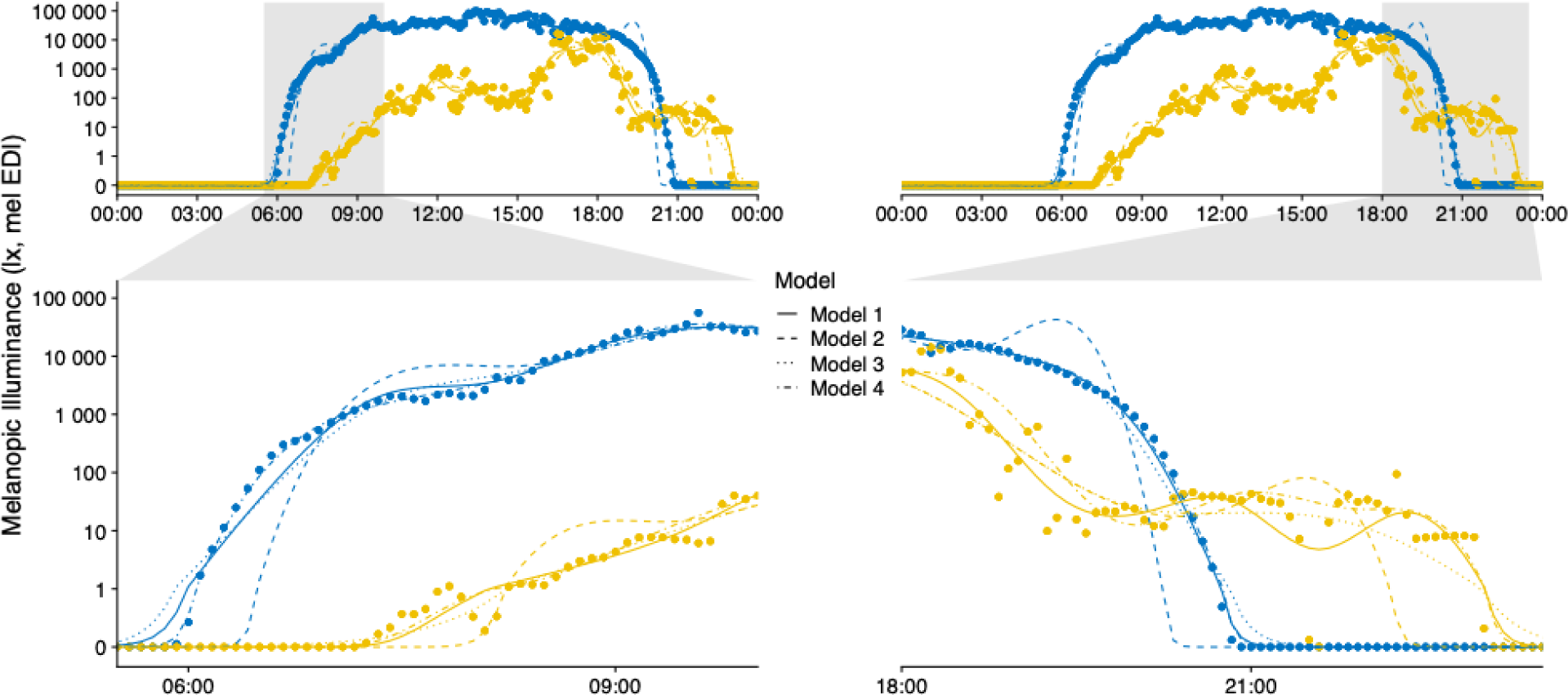
Combined Model results. Personal light exposure on a *symlog* scale for participant (yellow) and environmental light levels (blue). Points indicate measurements, lines indicate the fitted models. Zoomed facets show timeframes that cover transitions from and to zero-lux light levels.

## Discussion

### Visualizations of zero-inflated data

Figure 1 and Figure 2 show comparisons of typical (panels **A** and **B**) and atypical (panels **C** and **D**) ways to visualize light exposure patterns. While panel **A** shows impressive differences in light levels magnitude, it is inappropriate regarding sensation and perception of visual and non-visual systems [19]. Panel **B** is more appropriate in terms of the physiological receiving system. However, it overemphasizes values below 1 lux, which is neither sensible in the measurement accuracy of wearable devices nor in their relevance for the daily light exposure patterns. In other words, we would argue that a difference of 10^-2^ to 10^-3^ lux, is not as relevant as a difference of 10^2^ to 10^3^ lux in daily light exposure patterns, even if the light logger were to accurately measure <1 lux values. While the figure could be capped, e.g., at the level of measurement accuracy, it would still be ambiguous about whether the nighttime light levels were measured. Additionally, only absolute differences can be plotted (Figure 2B). Panel **C** removes this ambiguity but is harder to understand intuitively and inherits all other downsides mentioned for panel **B**. Panel **D** solves the previously mentioned issues by being clear to read and intuitive to understand, including zero-lux values, and without overemphasising small values. Further, it maps very well to showing heavy-tailed differences in both sign directions (Figure 2D). Beyond that, every point on the *symlog* scale can be mapped to a value by the viewer, as long as the threshold for logarithmic/linear transformation is provided, which is 1 lux in this case, but can be any value. This differentiates the *symlog* scale from, for example, the *pseudo-log* scale available in R [32]. *Pseudo-log* smoothes transitions between logarithmic and linear scale. While this approach allows for the same advantages that *symlog* provides, the smooth transition makes it harder to judge which scaling applies. We thus recommend the *symlog* scale as the preferred scaling for personal light exposure visualisations or similarly distributed measures.

### Statistical modelling of zero-inflated data

We modelled measures of personal light exposure on a logarithmic scale in four different ways. Specifically, we compared the model fits with the original training data to examine how dealing with zero-lux values influences model performance. While (generalized) additive models (GAMs) are not (yet) in common use for personal light exposure measurements, they are very well-fitted to deal with non-linear relationships typical in many natural and biological systems [33-35]. GAMs model the smooth relationship of the dependent variable to one or multiple linear or categorical predictors. Here, these are mel EDI as the dependent variable, and the time of day and the type of light exposure (environment or participant) as predictors. Further, compared to, for example, a generalized linear model, they not only allow comparison of model performance as a whole, but reveal where the model underperforms along the time series. These issues can be mapped to linear models, provided that they contain a similar composition of value ranges as the problematic areas in the additive model. In a standard analysis, data from more than one day or person would be modelled with a GAM in order to extract common trends or decompose the trend into contributing factors, such as chronotype or age. We limited our analysis to a single day to make the deviation of different models from the data directly visible.

Figure 3 reveals these problematic areas, especially pronounced for Model 2 (adding a very small value to the dataset to remove zero-lux values). These areas are transitional times, where the model must deal with zero-lux values and values of several orders of magnitude above in close proximity. Transitional times are shown in a larger zoom in Figure 4 to provide direct model comparisons. In the following paragraph, we first discuss each model individually and then draw comparisons between all models.

**Model 1** (removing zero-lux values, R^2^_adj_=0.949) captures the relevant daylight and participant features. The scaled residuals (Figure 3B) show very few patterns within their overall random dispersion. Model diagnostics, in general, are favourable (see analysis documentation **S1** section 4.1). The model performs quite well at zero-lux levels, considering it has no training data available in those instances. Transitional areas show only small deviations in the morning for environmental daylight levels. The greatest weakness of this model is that it is used to fit data outside of the (time) range of available data, which is generally not recommended [27]. Additionally, it is forced to disregard a substantial portion of actual and useful data that, from the model’s perspective, is indistinguishable from actual missing data.

**Model 2** (adding a very small non-zero value, R^2^_adj_=0.964) shows strong deviations from training data at the transitional times, which are obvious when comparing the model fit to the data (Figure 3A) and also the residuals (Figure 3B). The minuscule values at the original zero-lux level skew the logarithmically transformed value distribution. This leads to an overcompensation of the model fit on the other end of the distribution, manifesting as an overshoot. Other model diagnostics are also unfavourable, with strong patterns in the residuals (see analysis documentation **S1** section 4.2).

**Model 3** (adding -1 log_10_ lux, R^2^_adj_=0.981) captures the most relevant features of the daylight and participant data, with only slight deviations at transitional times. In the morning, this is visible in the direct comparison (Figure 3A), but residuals also reveal small structured deviations in the evening (Figure 3B). Other model diagnostics are good, with acceptable deviations from expected residual behaviour (see analysis documentation **S1** section 4.3).

**Model 4** (Tweedie error distribution, R^2^_adj_=0.917) balances pattern features in daylight and participant data and shows no deviations in the transitional periods or structured residual patterns. As a generalized model family, Tweedie is unsupported for the AR1 autoregressive error correction for additive models. As such, while other model diagnostics are good, there is a small amount of autocorrelation in the residuals (see analysis documentation **S1** section 4.4).

**In summary**, if zero-level values are removed (Model 1), model fitting is acceptable only in the scenario when few zero-lux instances exist, but not otherwise. Adding a common value across the dataset to make logarithmic values calculable (Model 2 and 3) skews logarithmic distributions and can lead to erroneous model fits. However, keeping a small distance on a logarithmic level dramatically reduces this effect, to the point where it performs better than when removing zero-level values outright (Model 3 vs. Model 1). We recommend using a value close to the lower bound of the manufacturer-specified measurement range. In our example, we added a -1 log_10_ lux value (0.1 lux) across the dataset for a lower measurement boundary of 0 log_10_ lux (1 lux). However, this procedure adds another step to data preparation and post-modelling, as model coefficients and/or predictions need to be readjusted back to the original value level. These steps can introduce new errors in analysis and reporting, especially for complex analyses. At a minimum, we recommend that the exact value added and the procedural steps must be unambiguously reported. Finally, a data transformation is not necessarily required, as the Tweedie error family includes zero values and models data with a logarithmic link function (Model 4).

While not the best model for R^2^, its performance is judged on the actual measurement data. This compares to Models 1 to 3, where R^2^_adj_ is based upon data that has already been transformed – and, in the case of Model 1, also pruned. Further, the adherence of Model 4 to the data during the transitional phases is remarkable (Figure 4). In terms of the features it captures, Model 4 also strikes a good balance between a slightly too *wiggly* [28] Model 1 (participant data), and the overall best-fit, but low on features Model 3. This is especially visible in the peaks throughout the day, which are slightly more pronounced in Model 4 than in Model 3 (Figure 3A). Thus, the Tweedie distribution is appropriate for modelling complex personal light exposure patterns on a logarithmic scale and including zero-lux values without any prior data transformation.

### Limitations

One avenue unexplored in this study is data aggregation until at least one value exceeds zero-lux. While certainly possible in some datasets, we find the loss of granularity to solve the issue of zero-lux values rather undesirable. The solution also does not generalize well, as different datasets would require a different level of aggregation. Some datasets, like the one presently used, would require the aggregation into 6-hour parts, thus eliminating most features.

Further, while our analysis systematically examines the consequences of different approaches, how it maps to other studies and analyses depends heavily on the statistical approach and distribution of data. However, it should hold strong for whole-day or longer periods of continuous data collection and (generalized) linear/additive (mixed) models.

Regarding visualizations, the recommended *symlog* transformation is seldom used to present and using it in scientific plots might seem unconventional to some. However, we believe its use is justified by the intuitive insight it allows, provided the scaling is explicitly stated, alongside the threshold where the transitions from linear to logarithmic scale occurs.

Finally, our approaches deal with small or zero-lux values “at face value”, i.e., not considering measurement errors. We believe that while values below the specified dynamic range of the wearable device might not be accurate to the same degree as values within, they contain valuable information about the overall light exposure pattern, and removing them outright would come at a loss for the bigger picture.

## Conclusion

Very small measurement values in light exposure patterns, especially zero, are regularly recorded in field studies with wearable devices. These values should neither be dismissed nor be unduly influential in visualizations and statistical models. Common types of visualization fail in at least one of those regards. We demonstrated that a *symlog*-transformed visualization style displays relevant features of light exposure across all scales, including zero-lux. Compared to pure logarithmic scaling, *symlog* reduces the emphasis on small values (e.g., <1 lux), which are less important in field studies that do not focus specifically on dark environments, like the sleep environment. Furthermore, this type of scaling maps very well to visualize differences in light exposure covering heavy-tailed negative values. The symlog scale is already part of the software package *LightLogR*, [24] and can thus be easily integrated into the analysis workflow.

We further showed that a small but not negligible value addition to the light exposure data for statistical modelling allows for acceptable models on a logarithmic scale. Those values should be close to the lower end of the measurement range on a logarithmic scale, such as 0.1 lux for a lower bound of 1 lux, as in our case. We also demonstrate the utility of the Tweedie distribution when modelling personal light exposure data, which does not require prior transformations, models data on a logarithmic scale, and includes zero-lux values. In conclusion, we provided a comprehensive analysis of the strengths and limitations of various approaches for addressing zero values in light exposure data. Our goal is to encourage researchers to approach zero-inflated datasets thoughtfully, make informed methodological choices, and ensure transparency in reporting their approaches.

## Data availability statement

Data are available on Zenodo [23] under a CC BY 4.0 license. The code is available as **Supplemental Material S1**.

## Contributions

Conceptualisation: JZ, CG, MS

Data curation: JZ, CG

Formal analysis: JZ

Funding acquisition: MS

Methodology: JZ

Project administration: JZ

Software: JZ

Resources: -

Supervision: -

Validation: -

Visualisation: JZ

Writing – original draft: JZ

Writing – review & editing: JZ, CG, MS

## Acknowledgements

None

## Supplementary Material

**S1. Analysis documentation.** HTML file that contains all code and code results to produce the output shown in the publication.

