## Supplement S1 for "How to deal with darkness: Modelling and visualization of zero-inflated personal light exposure data on a logarithmic scale"


#### Table of contents

- 1 Preface
- 2 Baseline Data and visualization
- 3 Visualizations
  - 3.1 Pattern
    - 3.1.1 Setup
    - 3.1.2 Visualization A
    - 3.1.3 Visualization B
    - 3.1.4 Visualization C
    - 3.1.5 Visualization D
    - 3.1.6 Combined Visualization
  - 3.2 Differences
    - 3.2.1 Visualization E
    - 3.2.2 Visualization F
    - 3.2.3 Visualization G
    - 3.2.4 Visualization H
    - 3.2.5 Combined Visualization
- 4 Modelling
  - 4.1 Model type 1: 0 to NA
  - 4.2 Model type 2: adding smallest float to every value
  - 4.3 Model type 3: adding -1 log10 to every value
  - 4.4 Model type 4: using the Tweedy family
  - 4.5 Combined Models
- 5 Session info

### Supplement S1

 Code

Author

Johannes Zauner

#### 1 Preface

this document contains the analysis documentation for the article **How to deal with darkness: Modelling and visualization of zero-inflated personal light exposure data on a logarithmic scale**.

```
# install.packages("devtools")
# devtools::install_github("tscnlab/LightLogR")

library(LightLogR)
library(tidyverse)
```

```
── Attaching core tidyverse packages ──────────────────────── tidyverse 2.0.0 ──
✔ dplyr     1.1.4     ✔ readr     2.1.5
✔ forcats   1.0.0     ✔ stringr   1.5.1
✔ ggplot2   3.5.1     ✔ tibble    3.2.1
✔ lubridate 1.9.4     ✔ tidyr     1.3.1
✔ purrr     1.0.2     
── Conflicts ────────────────────────────────────────── tidyverse_conflicts() ──
✖ dplyr::filter() masks stats::filter()
✖ dplyr::lag()    masks stats::lag()
ℹ Use the conflicted package (<http://conflicted.r-lib.org/>) to force all conflicts to become errors
```

```
library(mgcv)
```

```
Loading required package: nlme

Attaching package: 'nlme'

The following object is masked from 'package:dplyr':

    collapse

This is mgcv 1.9-1. For overview type 'help("mgcv-package")'.
```

```
library(patchwork)
library(gt)
library(itsadug)
```

```
Loading required package: plotfunctions

Attaching package: 'plotfunctions'

The following object is masked from 'package:ggplot2':

    alpha

Loaded package itsadug 2.4 (see 'help("itsadug")' ).
```

```
library(ggsci)
library(cowplot)
```

```
Attaching package: 'cowplot'

The following object is masked from 'package:gt':

    as_gtable

The following object is masked from 'package:patchwork':

    align_plots

The following object is masked from 'package:lubridate':

    stamp
```

```
library(ggforce)
```

#### 2 Baseline Data and visualization

In this section, we will copy relevant portions of a tutorial from LightLogR The whole game which will be the basis for both visualizations and analysis.

```
path <- system.file("extdata", 
              package = "LightLogR")

file.LL <- "205_actlumus_Log_1020_20230904101707532.txt.zip"
file.env <- "cyepiamb_CW35_Log_1431_20230904081953614.txt.zip"
tz <- "Europe/Berlin"
dataset.LL <- import$ActLumus(file.LL, path, auto.id = "^(\\d{3})", tz = tz)
```

```
Successfully read in 61'016 observations across 1 Ids from 1 ActLumus-file(s).
Timezone set is Europe/Berlin.

First Observation: 2023-08-28 08:47:54
Last Observation: 2023-09-04 10:17:04
Timespan: 7.1 days

Observation intervals: 
  Id    interval.time     n pct  
1 205   10s           61015 100%
```

```
dataset.env <- import$ActLumus(file.env, path, manual.id = "CW35", tz = tz)
```

```
Successfully read in 20'143 observations across 1 Ids from 1 ActLumus-file(s).
Timezone set is Europe/Berlin.

First Observation: 2023-08-28 08:28:39
Last Observation: 2023-09-04 08:19:38
Timespan: 7 days

Observation intervals: 
  Id    interval.time     n pct  
1 CW35  29s               1 0%   
2 CW35  30s           20141 100%
```

```
dataset.LL <- 
  dataset.LL %>% data2reference(Reference.data = dataset.env, across.id = TRUE)
dataset.LL <- 
  dataset.LL %>% select(Id, Datetime, MEDI, Reference)
dataset.LL.partial <- 
dataset.LL %>% #dataset
  filter_Date(start = "2023-09-01", length = days(1)) %>% 
  aggregate_Datetime(unit = "5 mins")
```

#### 3 Visualizations

##### 3.1 Pattern

###### 3.1.1 Setup

```
day_ribbon <- 
  geom_ribbon(aes(ymin = MEDI, ymax=Reference), 
              alpha = 0.25, fill = "#0073C2FF",
              outline.type = "upper", col = "#0073C2FF", size = 0.15)
```

```
Warning: Using `size` aesthetic for lines was deprecated in ggplot2 3.4.0.
ℹ Please use `linewidth` instead.
```

```
participant_ribbon <- 
  geom_ribbon(aes(ymin = 0, ymax = MEDI), alpha = 0.30, fill = "#EFC000", 
              outline.type = "upper", col = "#EFC000", size = 0.4)

lower_bound <- 
 geom_hline(yintercept = 1, col = "red", lty = 3)
upper_bound <- 
 geom_hline(yintercept = 10^5, col = "red", lty = 3)
time_scale <- 
  scale_x_time(breaks = c(0, 6, 12, 18, 24)*3600, labels = 
               scales::label_time(format = "%H:%M"), 
      expand = c(0, 0), limits = c(0, 24 * 3600))
```

###### 3.1.2 Visualization A

```
Plot_A <- 
dataset.LL.partial %>% 
   gg_day(facetting = FALSE, geom = "blank", y.scale = "identity",
          x.axis.label = "Local Time (HH:MM)",
          y.axis.breaks = seq(0, to = 10^5, by = 10^4), 
          y.axis.label = "Melanopic Illuminance (lx, mel EDI)") + #base plot
  day_ribbon + participant_ribbon +
 lower_bound + upper_bound + time_scale
```

```
Scale for x is already present.
Adding another scale for x, which will replace the existing scale.
```

```
Plot_A
```

###### 3.1.3 Visualization B

```
Plot_B <- 
dataset.LL.partial %>% 
   gg_day(facetting = FALSE, geom = "blank",
                    x.axis.label = "Local Time (HH:MM)",
          y.axis.label = "Melanopic Illuminance (lx, mel EDI)"
          ) + #base plot
  scale_y_log10(breaks = 10^(-5:5), labels = \(x) paste0("10^",log10(x))) +
  day_ribbon + participant_ribbon +
 lower_bound + upper_bound + time_scale
```

```
Scale for y is already present.
Adding another scale for y, which will replace the existing scale.
Scale for x is already present.
Adding another scale for x, which will replace the existing scale.
```

```
Plot_B
```

```
Warning in scale_y_log10(breaks = 10^(-5:5), labels = function(x) paste0("10^", : log-10 transformation introduced infinite values.
log-10 transformation introduced infinite values.
log-10 transformation introduced infinite values.
log-10 transformation introduced infinite values.
log-10 transformation introduced infinite values.
log-10 transformation introduced infinite values.
log-10 transformation introduced infinite values.
```

###### 3.1.4 Visualization C

```
Plot_C <-
dataset.LL.partial %>% 
  mutate(MEDI = case_when(MEDI == 0 ~ NA,
                          .default = MEDI),
         Reference = case_when(Reference == 0 ~ NA,
                               .default = Reference)) %>% 
   gg_day(facetting = FALSE, geom = "blank", 
                    x.axis.label = "Local Time (HH:MM)",
          y.axis.label = "Melanopic Illuminance (lx, mel EDI)"
          ) + #base plot
    geom_ribbon(
      aes(ymin = 
            ifelse(is.na(MEDI) | MEDI < min(Reference, na.rm = TRUE), 
                   min(Reference, na.rm = TRUE), MEDI), 
          ymax=Reference), 
              alpha = 0.25, fill = "#0073C2FF",
              outline.type = "upper", col = "#0073C2FF", size = 0.15) + #solar reference
  geom_ribbon(aes(ymin = min(MEDI, na.rm = TRUE), ymax = MEDI), alpha = 0.30, 
              fill = "#EFC000",
              outline.type = "upper", col = "#EFC000", size = 0.4) +
  geom_point(
    data = \(x) x %>% filter(is.na(Reference)) %>% mutate(Reference = 10^-4), 
    aes(y = Reference), col = "#0073C2FF", size = 1, alpha = 0.25) +
  geom_point(data = \(x) x %>% filter(is.na(MEDI)) %>% mutate(MEDI = 10^-4), 
              col = "#EFC000", size = 1, alpha = 0.25, position = position_nudge(y=0.05)) +
  scale_y_log10(breaks = 10^(-5:5), 
                labels = \(x) ifelse(x == 10^-4, "0", paste0("10^",log10(x)))
                ) +
 lower_bound + upper_bound + time_scale
```

```
Scale for y is already present.
Adding another scale for y, which will replace the existing scale.
Scale for x is already present.
Adding another scale for x, which will replace the existing scale.
```

```
Plot_C
```

###### 3.1.5 Visualization D

```
Plot_D <-
dataset.LL.partial %>% 
   gg_day(facetting = FALSE, geom = "blank", 
                    x.axis.label = "Local Time (HH:MM)",
          y.axis.label = "Melanopic Illuminance (lx, mel EDI)") + #base plot
 day_ribbon + participant_ribbon +
 lower_bound + upper_bound + time_scale
```

```
Scale for x is already present.
Adding another scale for x, which will replace the existing scale.
```

```
Plot_D
```

###### 3.1.6 Combined Visualization

```
Pattern <-
Plot_A + plot_spacer() + Plot_B + plot_spacer() + Plot_C + plot_spacer() + Plot_D + 
  plot_layout(ncol = 7, nrow = 1, axes = "collect",
              widths = c(1,-0.15,1,-0.15,1,-0.15,1)) + 
  plot_annotation(tag_levels = "A") & 
  theme(
    plot.tag.position = c(0, 1),
        plot.tag.location = "plot",
        plot.tag = element_text(size = 15, hjust = 0.5, vjust =-0.75),
        axis.text = element_text(size = 13),
        axis.title = element_text(size = 16)
        ) +
  theme(plot.margin = margin(15,10,5,5))

ggsave("figures/Figure_1.pdf", Pattern, width = 17.5, height = 5.5, units = "cm", scale = 2)
```

```
Warning in scale_y_log10(breaks = 10^(-5:5), labels = function(x) paste0("10^", : log-10 transformation introduced infinite values.
log-10 transformation introduced infinite values.
log-10 transformation introduced infinite values.
log-10 transformation introduced infinite values.
log-10 transformation introduced infinite values.
log-10 transformation introduced infinite values.
log-10 transformation introduced infinite values.
```

##### 3.2 Differences

###### 3.2.1 Visualization E

```
Plot_E <-
dataset.LL.partial %>% 
   gg_day(y.axis = Reference - MEDI, facetting = FALSE, geom = "blank", 
                    x.axis.label = "Local Time (HH:MM)", y.scale = "identity",
          y.axis.breaks = seq(-10^4, to = 10^5, by = 10^4),
          y.axis.label = "Daylight - Participant (lx, mel EDI)") + #base plot
 geom_area(
   aes(group = consecutive_id((Reference - MEDI) >=0 ),
       fill = (Reference - MEDI) >=0,
       col = (Reference - MEDI) >=0
       ), outline.type = "upper", alpha = 0.25, size = 0.25
 )+
  guides(fill = "none", col = "none") +
  time_scale +
  geom_hline(aes(yintercept = 0), lty = 2, col = "red") + 
    scale_fill_manual(values = c("#EFC000", "#0073C2")) + 
    scale_color_manual(values = c("#EFC000", "#0073C2"))
```

```
Scale for x is already present.
Adding another scale for x, which will replace the existing scale.
Scale for fill is already present.
Adding another scale for fill, which will replace the existing scale.
Scale for colour is already present.
Adding another scale for colour, which will replace the existing scale.
```

```
Plot_E
```

###### 3.2.2 Visualization F

```
Plot_F <-
dataset.LL.partial %>% 
   gg_day(facetting = FALSE, geom = "blank", 
                    x.axis.label = "Local Time (HH:MM)",
          y.axis.label = "Daylight - Participant (lx, mel EDI)") + #base plot
 geom_ribbon(
   aes(
     ymin = 0, ymax = abs(Reference - MEDI),
     group = consecutive_id((Reference - MEDI) >=0 ),
       fill = (Reference - MEDI) >=0,
       col = (Reference - MEDI) >=0
       ), outline.type = "upper", alpha = 0.25, size = 0.25
 )+
  guides(fill = "none", col = "none") +
  time_scale +
  geom_hline(aes(yintercept = 0), lty = 2, col = "red") + 
    scale_fill_manual(values = c("#EFC000", "#0073C2")) + 
    scale_color_manual(values = c("#EFC000", "#0073C2")) +
    scale_y_log10(breaks = 10^(-5:5), labels = \(x) paste0("10^",log10(x)))
```

```
Scale for x is already present.
Adding another scale for x, which will replace the existing scale.
Scale for fill is already present.
Adding another scale for fill, which will replace the existing scale.
Scale for colour is already present.
Adding another scale for colour, which will replace the existing scale.
Scale for y is already present.
Adding another scale for y, which will replace the existing scale.
```

```
Plot_F
```

```
Warning in scale_y_log10(breaks = 10^(-5:5), labels = function(x) paste0("10^", : log-10 transformation introduced infinite values.
log-10 transformation introduced infinite values.
log-10 transformation introduced infinite values.
log-10 transformation introduced infinite values.
log-10 transformation introduced infinite values.
```

###### 3.2.3 Visualization G

```
Plot_G <-
dataset.LL.partial %>% 
  mutate(MEDI = ifelse(Reference-MEDI == 0, NA, MEDI)) %>% 
   gg_day(facetting = FALSE, geom = "blank", 
                    x.axis.label = "Local Time (HH:MM)",
          y.axis.label = "Daylight - Participant (lx, mel EDI)") + #base plot
 geom_ribbon(
   aes(
     ymin = min(abs(Reference-MEDI), na.rm = TRUE), ymax = abs(Reference - MEDI),
     group = consecutive_id((Reference - MEDI) >=0 ),
       fill = (Reference - MEDI) >=0,
       col = (Reference - MEDI) >=0
       ), outline.type = "upper", alpha = 0.25, size = 0.25
 )+
  guides(fill = "none", col = "none") +
  time_scale +
  geom_hline(aes(yintercept = 10^-4), lty = 2, col = "red") + 
    scale_fill_manual(values = c("#EFC000", "#0073C2")) + 
    scale_color_manual(values = c("#EFC000", "#0073C2")) +
  scale_y_log10(breaks = 10^(-5:5), 
                labels = \(x) ifelse(x == 10^-4, "0", paste0("10^",log10(x)))
                ) +
  geom_point(
    data = \(x) x %>% filter(is.na(MEDI)) %>% mutate(MEDI = 10^-4), 
    aes(y = MEDI), col = "grey", size = 1, alpha = 0.5)
```

```
Scale for x is already present.
Adding another scale for x, which will replace the existing scale.
Scale for fill is already present.
Adding another scale for fill, which will replace the existing scale.
Scale for colour is already present.
Adding another scale for colour, which will replace the existing scale.
Scale for y is already present.
Adding another scale for y, which will replace the existing scale.
```

```
Plot_G
```

```
Warning in scale_y_log10(breaks = 10^(-5:5), labels = function(x) ifelse(x == : log-10 transformation introduced infinite values.
log-10 transformation introduced infinite values.
```

```
Warning in max(ids, na.rm = TRUE): no non-missing arguments to max; returning
-Inf
Warning in max(ids, na.rm = TRUE): no non-missing arguments to max; returning
-Inf
```

###### 3.2.4 Visualization H

```
Plot_H <-
dataset.LL.partial %>% 
   gg_day(y.axis = Reference - MEDI, facetting = FALSE, geom = "blank", 
                    x.axis.label = "Local Time (HH:MM)",
          y.axis.label = "Daylight - Participant (lx, mel EDI)") + #base plot
 geom_area(
   aes(group = consecutive_id((Reference - MEDI) >=0 ),
       fill = (Reference - MEDI) >=0,
       col = (Reference - MEDI) >=0
       ), outline.type = "upper", alpha = 0.25, size = 0.25
 )+
  guides(fill = "none", col = "none") +
  time_scale +
  geom_hline(aes(yintercept = 0), lty = 2, col = "red") + 
    scale_fill_manual(values = c("#EFC000", "#0073C2")) + 
    scale_color_manual(values = c("#EFC000", "#0073C2"))
```

```
Scale for x is already present.
Adding another scale for x, which will replace the existing scale.
Scale for fill is already present.
Adding another scale for fill, which will replace the existing scale.
Scale for colour is already present.
Adding another scale for colour, which will replace the existing scale.
```

```
Plot_H
```

###### 3.2.5 Combined Visualization

```
Differences <- 
Plot_E + plot_spacer() + Plot_F + plot_spacer() + Plot_G + plot_spacer() + Plot_H + 
  plot_layout(ncol = 7, nrow = 1, axes = "collect",
              widths = c(1,-0.15,1,-0.15,1,-0.15,1)) + 
  plot_annotation(tag_levels = "A") & 
  theme(
    plot.tag.position = c(0, 1),
        plot.tag.location = "plot",
        plot.tag = element_text(size = 15, hjust = 0, vjust =-0.75),
        axis.text = element_text(size = 13),
        axis.title = element_text(size = 16)
        ) +
  theme(plot.margin = margin(15,10,5,5))

ggsave("figures/Figure_2.pdf", Differences, width = 17.5, height = 5.5, units = "cm", scale = 2)
```

```
Warning in scale_y_log10(breaks = 10^(-5:5), labels = function(x) paste0("10^", : log-10 transformation introduced infinite values.
log-10 transformation introduced infinite values.
log-10 transformation introduced infinite values.
log-10 transformation introduced infinite values.
log-10 transformation introduced infinite values.
```

```
Warning in scale_y_log10(breaks = 10^(-5:5), labels = function(x) ifelse(x == : log-10 transformation introduced infinite values.
log-10 transformation introduced infinite values.
```

```
Warning in max(ids, na.rm = TRUE): no non-missing arguments to max; returning
-Inf
Warning in max(ids, na.rm = TRUE): no non-missing arguments to max; returning
-Inf
```

#### 4 Modelling

```
#setting the ends for the cyclic smooth
knots_day <- list(time = c(0, 24*3600))

model_data <- 
dataset.LL.partial %>% 
  ungroup() %>% 
  rename(Environment = Reference) %>% 
  pivot_longer(names_to = "type", cols = c(MEDI, Environment)) %>%  
  arrange(type) %>% 
  mutate(time = hms::as_hms(Datetime) %>% as.numeric(),
         time = c(time[-n()], 24*3600),
         type = factor(type),
         start.event = time == 0,
         input_m1 = 
           case_when(
             value == 0 ~ NA,
             .default = value),
         input_m2 = value + .Machine$double.eps,
         input_m3 = value + 0.1,
         input_m4 = value,
         .by = type
  )
```

##### 4.1 Model type 1: 0 to NA

```
pattern_formula <- log10(input_m1) ~ type + s(time, by = type, bs = "cc", k = 24)

model_data_m1 <- 
  model_data %>% 
  group_by(type, case = is.na(input_m1)) %>% 
  mutate(start.event = 
                        c(TRUE, rep(FALSE, n() -1)))

model_1 <- bam(formula = pattern_formula, 
             data = model_data_m1, 
             method = "REML", 
             knots = knots_day)

r1 <- start_value_rho(model_1, plot=TRUE)
```

```
model_1 <- bam(formula = pattern_formula, 
             data = model_data_m1, 
             method = "REML", 
             knots = knots_day, 
             rho = r1,
             AR.start = model_data_m1$start.event
              )

acf_resid(model_1)
```

```
summary(model_1)
```

```
Family: gaussian 
Link function: identity 

Formula:
log10(input_m1) ~ type + s(time, by = type, bs = "cc", k = 24)

Parametric coefficients:
            Estimate Std. Error t value Pr(>|t|)    
(Intercept)  3.37020    0.09289   36.28   <2e-16 ***
typeMEDI    -1.81237    0.10998  -16.48   <2e-16 ***
---
Signif. codes:  0 '***' 0.001 '**' 0.01 '*' 0.05 '.' 0.1 ' ' 1

Approximate significance of smooth terms:
                          edf Ref.df     F p-value    
s(time):typeEnvironment 14.45     22 43.18  <2e-16 ***
s(time):typeMEDI        18.49     22 42.94  <2e-16 ***
---
Signif. codes:  0 '***' 0.001 '**' 0.01 '*' 0.05 '.' 0.1 ' ' 1

R-sq.(adj) =  0.949   Deviance explained = 95.3%
-REML = 189.85  Scale est. = 0.12884   n = 371
```

```
gam.check(model_1, rep = 500)
```

```
Method: REML   Optimizer: outer newton
full convergence after 6 iterations.
Gradient range [-1.632827e-10,8.787637e-12]
(score 189.8507 & scale 0.1288363).
Hessian positive definite, eigenvalue range [2.029136,185.2563].
Model rank =  46 / 46 

Basis dimension (k) checking results. Low p-value (k-index<1) may
indicate that k is too low, especially if edf is close to k'.

                          k'  edf k-index p-value   
s(time):typeEnvironment 22.0 14.5    0.87   0.015 * 
s(time):typeMEDI        22.0 18.5    0.87   0.010 **
---
Signif. codes:  0 '***' 0.001 '**' 0.01 '*' 0.05 '.' 0.1 ' ' 1
```

```
estimate_m1 <- 
plot_smooth(model_1, 
            view = "time", 
            plot_all = "type",
            # rug = F,
            n.grid = 12*24+1,
            rm.ranef = FALSE,
            se = 0)
```

```
Summary:
    * type : factor; set to the value(s): Environment, MEDI. 
    * time : numeric predictor; with 289 values ranging from 0.000000 to 86400.000000.
```

```
estimate_m1_dif <- 
plot_diff(model_1,
          view = "time",
          comp = list(type = c("Environment", "MEDI")),
          rm.ranef = FALSE,
          n.grid = 12*24+1)
```

```
Summary:
    * time : numeric predictor; with 289 values ranging from 21300.000000 to 82800.000000.
```

```
time window(s) of significant difference(s):
    21300.000000 - 72977.083333
    73831.250000 - 82800.000000
```

```
estimate_m1$fv %>% 
  ggplot(aes(x = time, y = 10^fit, col = type)) +
  geom_line() +
  theme_minimal() +
  scale_x_time(limits = c(0, 24*60*60), expand = c(0,0),
               breaks = c(0, 6, 12, 18, 24)*60*60, 
               labels = scales::label_time(format = "%H:%M")
               ) +
  scale_y_continuous(transform = "symlog", breaks = c(0, 10^(0:5))) +
  geom_point(data = model_data, aes(y = value, col = type))
```

```
model_data <- 
model_data %>% 
  left_join(estimate_m1$fv, by = c("type", "time")) %>% 
  rename(fit_m1 = fit) %>% 
  mutate(fit_m1 = (10^(fit_m1)))
```

##### 4.2 Model type 2: adding smallest float to every value

```
pattern_formula <- log10(input_m2) ~ type + s(time, by = type, bs = "cc", k = 24)

model_2 <- bam(formula = pattern_formula, 
             data = model_data, 
             method = "REML", 
             knots = knots_day)

r1 <- start_value_rho(model_2, plot=TRUE)
```

```
model_2 <- bam(formula = pattern_formula, 
             data = model_data, 
             method = "REML", 
             knots = knots_day, 
             rho = r1,
             AR.start = model_data$start.event
              )

acf_resid(model_2)
```

```
summary(model_2)
```

```
Family: gaussian 
Link function: identity 

Formula:
log10(input_m2) ~ type + s(time, by = type, bs = "cc", k = 24)

Parametric coefficients:
            Estimate Std. Error t value Pr(>|t|)    
(Intercept)  -3.4604     0.2267 -15.262   <2e-16 ***
typeMEDI     -0.6504     0.3206  -2.029    0.043 *  
---
Signif. codes:  0 '***' 0.001 '**' 0.01 '*' 0.05 '.' 0.1 ' ' 1

Approximate significance of smooth terms:
                          edf Ref.df     F p-value    
s(time):typeEnvironment 16.83     22 79.67  <2e-16 ***
s(time):typeMEDI        15.60     22 59.99  <2e-16 ***
---
Signif. codes:  0 '***' 0.001 '**' 0.01 '*' 0.05 '.' 0.1 ' ' 1

R-sq.(adj) =  0.964   Deviance explained = 96.6%
-REML = 949.28  Scale est. = 2.5616    n = 578
```

```
gam.check(model_2, rep = 500)
```

```
Method: REML   Optimizer: outer newton
full convergence after 5 iterations.
Gradient range [-1.510424e-07,4.404574e-09]
(score 949.2801 & scale 2.561581).
Hessian positive definite, eigenvalue range [4.66738,288.4646].
Model rank =  46 / 46 

Basis dimension (k) checking results. Low p-value (k-index<1) may
indicate that k is too low, especially if edf is close to k'.

                          k'  edf k-index p-value
s(time):typeEnvironment 22.0 16.8    0.99    0.46
s(time):typeMEDI        22.0 15.6    0.99    0.48
```

```
estimate_m2 <- 
plot_smooth(model_2, 
            view = "time", 
            plot_all = "type",
            # rug = F,
            n.grid = 12*24+1,
            rm.ranef = FALSE,
            se = 0)
```

```
Summary:
    * type : factor; set to the value(s): Environment, MEDI. 
    * time : numeric predictor; with 289 values ranging from 0.000000 to 86400.000000.
```

```
estimate_m2_dif <- 
plot_diff(model_2,
          view = "time",
          comp = list(type = c("Environment", "MEDI")),
          rm.ranef = FALSE,
          n.grid = 12*24+1)
```

```
Summary:
    * time : numeric predictor; with 289 values ranging from 0.000000 to 86400.000000.
```

```
time window(s) of significant difference(s):
    19500.000000 - 39900.000000
    47100.000000 - 55200.000000
    69300.000000 - 70500.000000
    73500.000000 - 84900.000000
```

```
estimate_m2$fv %>% 
  ggplot(aes(x = time, y = (10^fit)-.Machine$double.eps, col = type)) +
  geom_line() +
  theme_minimal() +
  scale_x_time(limits = c(0, 24*60*60), expand = c(0,0),
               breaks = c(0, 6, 12, 18, 24)*60*60, 
               labels = scales::label_time(format = "%H:%M")
               ) +
  scale_y_continuous(transform = "symlog", breaks = c(0, 10^(0:5))) +
  geom_point(data = model_data, aes(y = value, col = type))
```

```
model_data <- 
model_data %>% 
  left_join(estimate_m2$fv, by = c("type", "time")) %>% 
  rename(fit_m2 = fit) %>% 
  mutate(fit_m2 = (10^(fit_m2)) - .Machine$double.eps)
```

##### 4.3 Model type 3: adding -1 log10 to every value

```
pattern_formula <- log10(input_m3) ~ type + s(time, by = type, bs = "cc", k = 24)

model_3 <- bam(formula = pattern_formula, 
             data = model_data, 
             method = "REML", 
             knots = knots_day)

r1 <- start_value_rho(model_3, plot=TRUE)
```

```
model_3 <- bam(formula = pattern_formula,
             data = model_data,
             method = "REML",
             knots = knots_day,
             rho = r1,
             AR.start = model_data$start.event
              )

acf_resid(model_3)
```

```
summary(model_3)
```

```
Family: gaussian 
Link function: identity 

Formula:
log10(input_m3) ~ type + s(time, by = type, bs = "cc", k = 24)

Parametric coefficients:
            Estimate Std. Error t value Pr(>|t|)    
(Intercept)  2.06937    0.03919   52.80   <2e-16 ***
typeMEDI    -1.19000    0.05542  -21.47   <2e-16 ***
---
Signif. codes:  0 '***' 0.001 '**' 0.01 '*' 0.05 '.' 0.1 ' ' 1

Approximate significance of smooth terms:
                          edf Ref.df      F p-value    
s(time):typeEnvironment 17.62     22 194.49  <2e-16 ***
s(time):typeMEDI        14.48     22  75.97  <2e-16 ***
---
Signif. codes:  0 '***' 0.001 '**' 0.01 '*' 0.05 '.' 0.1 ' ' 1

R-sq.(adj) =  0.981   Deviance explained = 98.2%
-REML = 21.691  Scale est. = 0.090725  n = 578
```

```
gam.check(model_3, rep = 500)
```

```
Method: REML   Optimizer: outer newton
full convergence after 5 iterations.
Gradient range [-1.22994e-07,1.919258e-09]
(score 21.69085 & scale 0.09072546).
Hessian positive definite, eigenvalue range [5.504781,288.4603].
Model rank =  46 / 46 

Basis dimension (k) checking results. Low p-value (k-index<1) may
indicate that k is too low, especially if edf is close to k'.

                          k'  edf k-index p-value
s(time):typeEnvironment 22.0 17.6     1.1    0.99
s(time):typeMEDI        22.0 14.5     1.1    1.00
```

```
estimate_m3 <- 
plot_smooth(model_3, 
            view = "time", 
            plot_all = "type",
            # rug = F,
            n.grid = 12*24+1,
            rm.ranef = FALSE,
            se = 0)
```

```
Summary:
    * type : factor; set to the value(s): Environment, MEDI. 
    * time : numeric predictor; with 289 values ranging from 0.000000 to 86400.000000.
```

```
estimate_m3_dif <- 
plot_diff(model_3,
          view = "time",
          comp = list(type = c("Environment", "MEDI")),
          rm.ranef = FALSE,
          n.grid = 12*24+1)
```

```
Summary:
    * time : numeric predictor; with 289 values ranging from 0.000000 to 86400.000000.
```

```
time window(s) of significant difference(s):
    20100.000000 - 72900.000000
    74100.000000 - 84600.000000
```

```
estimate_m3$fv %>% 
  ggplot(aes(x = time, y = (10^fit)-0.1, col = type)) +
  geom_line() +
  theme_minimal() +
  scale_x_time(limits = c(0, 24*60*60), expand = c(0,0),
               breaks = c(0, 6, 12, 18, 24)*60*60, 
               labels = scales::label_time(format = "%H:%M")
               ) +
  scale_y_continuous(transform = "symlog", breaks = c(0, 10^(0:5))) +
  geom_point(data = model_data, aes(y = value, col = type))
```

```
model_data <- 
model_data %>% 
  left_join(estimate_m3$fv, by = c("type", "time")) %>% 
  rename(fit_m3 = fit) %>% 
  mutate(fit_m3 = (10^(fit_m3))-0.1)
```

##### 4.4 Model type 4: using the Tweedy family

```
pattern_formula <- input_m4 ~ type + s(time, by = type, bs = "cc", k = 24)

model_4 <- gam(formula = pattern_formula, 
             data = model_data, 
             method = "REML", 
             family = tw,
             knots = knots_day)

summary(model_4)
```

```
Family: Tweedie(p=1.726) 
Link function: log 

Formula:
input_m4 ~ type + s(time, by = type, bs = "cc", k = 24)

Parametric coefficients:
            Estimate Std. Error t value Pr(>|t|)   
(Intercept)   -4.654      1.704  -2.732  0.00651 **
typeMEDI      -0.247      2.173  -0.114  0.90957   
---
Signif. codes:  0 '***' 0.001 '**' 0.01 '*' 0.05 '.' 0.1 ' ' 1

Approximate significance of smooth terms:
                          edf Ref.df     F p-value    
s(time):typeEnvironment 15.66     22 119.3  <2e-16 ***
s(time):typeMEDI        16.61     22 114.7  <2e-16 ***
---
Signif. codes:  0 '***' 0.001 '**' 0.01 '*' 0.05 '.' 0.1 ' ' 1

R-sq.(adj) =  0.917   Deviance explained = 97.8%
-REML = 2895.9  Scale est. = 2.0844    n = 578
```

```
gam.check(model_4, rep = 500)
```

```
Method: REML   Optimizer: outer newton
full convergence after 5 iterations.
Gradient range [-0.005965489,0.0009400968]
(score 2895.865 & scale 2.084384).
Hessian positive definite, eigenvalue range [2.112661,585.8741].
Model rank =  46 / 46 

Basis dimension (k) checking results. Low p-value (k-index<1) may
indicate that k is too low, especially if edf is close to k'.

                          k'  edf k-index p-value
s(time):typeEnvironment 22.0 15.7    1.12       1
s(time):typeMEDI        22.0 16.6    1.12       1
```

```
plot(model_4)
```

```
estimate_m4 <- 
plot_smooth(model_4, 
            view = "time", 
            plot_all = "type",
            # rug = F,
            n.grid = 12*24+1,
            rm.ranef = FALSE,
            se = 0)
```

```
Summary:
    * type : factor; set to the value(s): Environment, MEDI. 
    * time : numeric predictor; with 289 values ranging from 0.000000 to 86400.000000.
```

```
estimate_m4_dif <- 
plot_diff(model_4,
          view = "time",
          comp = list(type = c("Environment", "MEDI")),
          rm.ranef = FALSE,
          n.grid = 12*24+1)
```

```
Summary:
    * time : numeric predictor; with 289 values ranging from 0.000000 to 86400.000000.
```

```
time window(s) of significant difference(s):
    0.000000 - 1500.000000
    18300.000000 - 73200.000000
    74100.000000 - 86400.000000
```

```
estimate_m4$fv %>% 
  ggplot(aes(x = time, y = exp(fit), col = type)) +
  geom_line() +
  theme_minimal() +
  scale_x_time(limits = c(0, 24*60*60), expand = c(0,0),
               breaks = c(0, 6, 12, 18, 24)*60*60, 
               labels = scales::label_time(format = "%H:%M")
               ) +
  scale_y_continuous(transform = "symlog", breaks = c(0, 10^(0:5))) +
  geom_point(data = model_data, aes(y = value, col = type))
```

```
model_data <- 
model_data %>% 
  left_join(estimate_m4$fv, by = c("type", "time")) %>% 
  rename(fit_m4 = fit) %>% 
  mutate(fit_m4 = exp(fit_m4))
```

##### 4.5 Combined Models

```
#adding scaled.pearson residuals
model_data$resid_m1 <- NA
model_data$resid_m1[!is.na(model_data$input_m1)] <- 
  (residuals(model_1, type = "scaled.pearson"))
model_data$resid_m2 <-(residuals(model_2, type = "scaled.pearson"))
model_data$resid_m3 <- (residuals(model_3, type = "scaled.pearson"))
model_data$resid_m4 <- (residuals(model_4, type = "scaled.pearson"))

model_data_long <- 
model_data %>% 
  pivot_longer(cols = input_m1:resid_m4, names_to = c(".value", "model"), 
               names_pattern = "(.*)_(m\\d{1})") %>% 
  mutate(model = model %>% str_replace(pattern = "m", replacement =  "Model "))


summary(model_1)
```

```
Family: gaussian 
Link function: identity 

Formula:
log10(input_m1) ~ type + s(time, by = type, bs = "cc", k = 24)

Parametric coefficients:
            Estimate Std. Error t value Pr(>|t|)    
(Intercept)  3.37020    0.09289   36.28   <2e-16 ***
typeMEDI    -1.81237    0.10998  -16.48   <2e-16 ***
---
Signif. codes:  0 '***' 0.001 '**' 0.01 '*' 0.05 '.' 0.1 ' ' 1

Approximate significance of smooth terms:
                          edf Ref.df     F p-value    
s(time):typeEnvironment 14.45     22 43.18  <2e-16 ***
s(time):typeMEDI        18.49     22 42.94  <2e-16 ***
---
Signif. codes:  0 '***' 0.001 '**' 0.01 '*' 0.05 '.' 0.1 ' ' 1

R-sq.(adj) =  0.949   Deviance explained = 95.3%
-REML = 189.85  Scale est. = 0.12884   n = 371
```

```
summary(model_2)
```

```
Family: gaussian 
Link function: identity 

Formula:
log10(input_m2) ~ type + s(time, by = type, bs = "cc", k = 24)

Parametric coefficients:
            Estimate Std. Error t value Pr(>|t|)    
(Intercept)  -3.4604     0.2267 -15.262   <2e-16 ***
typeMEDI     -0.6504     0.3206  -2.029    0.043 *  
---
Signif. codes:  0 '***' 0.001 '**' 0.01 '*' 0.05 '.' 0.1 ' ' 1

Approximate significance of smooth terms:
                          edf Ref.df     F p-value    
s(time):typeEnvironment 16.83     22 79.67  <2e-16 ***
s(time):typeMEDI        15.60     22 59.99  <2e-16 ***
---
Signif. codes:  0 '***' 0.001 '**' 0.01 '*' 0.05 '.' 0.1 ' ' 1

R-sq.(adj) =  0.964   Deviance explained = 96.6%
-REML = 949.28  Scale est. = 2.5616    n = 578
```

```
summary(model_3)
```

```
Family: gaussian 
Link function: identity 

Formula:
log10(input_m3) ~ type + s(time, by = type, bs = "cc", k = 24)

Parametric coefficients:
            Estimate Std. Error t value Pr(>|t|)    
(Intercept)  2.06937    0.03919   52.80   <2e-16 ***
typeMEDI    -1.19000    0.05542  -21.47   <2e-16 ***
---
Signif. codes:  0 '***' 0.001 '**' 0.01 '*' 0.05 '.' 0.1 ' ' 1

Approximate significance of smooth terms:
                          edf Ref.df      F p-value    
s(time):typeEnvironment 17.62     22 194.49  <2e-16 ***
s(time):typeMEDI        14.48     22  75.97  <2e-16 ***
---
Signif. codes:  0 '***' 0.001 '**' 0.01 '*' 0.05 '.' 0.1 ' ' 1

R-sq.(adj) =  0.981   Deviance explained = 98.2%
-REML = 21.691  Scale est. = 0.090725  n = 578
```

```
summary(model_4)
```

```
Family: Tweedie(p=1.726) 
Link function: log 

Formula:
input_m4 ~ type + s(time, by = type, bs = "cc", k = 24)

Parametric coefficients:
            Estimate Std. Error t value Pr(>|t|)   
(Intercept)   -4.654      1.704  -2.732  0.00651 **
typeMEDI      -0.247      2.173  -0.114  0.90957   
---
Signif. codes:  0 '***' 0.001 '**' 0.01 '*' 0.05 '.' 0.1 ' ' 1

Approximate significance of smooth terms:
                          edf Ref.df     F p-value    
s(time):typeEnvironment 15.66     22 119.3  <2e-16 ***
s(time):typeMEDI        16.61     22 114.7  <2e-16 ***
---
Signif. codes:  0 '***' 0.001 '**' 0.01 '*' 0.05 '.' 0.1 ' ' 1

R-sq.(adj) =  0.917   Deviance explained = 97.8%
-REML = 2895.9  Scale est. = 2.0844    n = 578
```

```
#model fitted plot
Model_fitted_plot <- 
model_data_long %>%
  ggplot(aes(x = time, y = value, group = type, col = type)) +
  geom_point(size = 0.5) +
  geom_line(aes(y = fit), col = "black") +
  facet_wrap(~model, ncol = 4) + 
  # geom_line(col = "red") +
  # geom_line(aes(y = fit_m2), col = "skyblue3") +
  # geom_line(aes(y = fit_m3), col = "limegreen") +
  # geom_line(aes(y = fit_m4), col = "orange2") +
  theme_cowplot() +
  scale_x_time(limits = c(0, 24*60*60), expand = c(0,0),
               breaks = c(0, 6, 12, 18, 24)*60*60, 
               labels = scales::label_time(format = "%H:%M")
               ) +
  scale_y_continuous(transform = "symlog", breaks = c(0, 10^(0:5)),
                     labels = c("0", "1", "10", "100", "1 000", "10 000", "100 000")) +
  scale_color_jco() +
  guides(color = "none") +
  labs(y = "Melanopic Illuminance\n(lx, mel EDI)",
       x = NULL) +
  theme(
        strip.text = element_text(size = 14)
  )
 

Model_residual_plot <- 
model_data_long %>%   
ggplot(aes(x=time))+
  geom_point(aes(y=resid), size = 0.5) +
  facet_wrap(~model, ncol = 4) +
  theme_cowplot() +
  coord_cartesian(ylim = c(-5,5)) +
  time_scale +
  labs(y = "Scaled Pearson\nresiduals", x = "Local time (HH:MM)") +
  theme(strip.background = element_blank(),
        strip.text = element_blank())

Model_fitted_plot / Model_residual_plot + 
  plot_layout(guides = "collect", heights = c(3,2)) +
  plot_annotation(tag_levels = "A") &
  theme(
    plot.tag.position = c(0, 0.99),
    plot.tag = element_text(size = 15, hjust = 0.5, vjust = 0, face = "bold"),
    panel.spacing = unit(2.5, "lines"),
        plot.tag.location = "plot",
        axis.text = element_text(size = 13),
        axis.title = element_text(size = 16)
        ) +
  theme(plot.margin = margin(15,10,5,5))
```

```
Warning: Removed 207 rows containing missing values or values outside the scale range
(`geom_point()`).
```

```
  ggsave("figures/Figure_3.pdf", width = 17.5, height = 8, units = "cm", scale = 2)
```

```
Warning: Removed 207 rows containing missing values or values outside the scale range
(`geom_point()`).
```

```
Model_zoom1 <- 
model_data_long %>%
  ggplot(aes(x = time, y = value, group = type, col = type)) +
  geom_point(size = 1.5) +
  geom_line(aes(y = fit)) +
  facet_wrap(~model, ncol = 4) + 
  # geom_line(col = "red") +
  # geom_line(aes(y = fit_m2), col = "skyblue3") +
  # geom_line(aes(y = fit_m3), col = "limegreen") +
  # geom_line(aes(y = fit_m4), col = "orange2") +
  theme_cowplot() +
  scale_x_time(limits = c(0, 24*60*60), expand = c(0,0),
               breaks = c(0, 3, 6, 9, 12, 15, 18, 21, 24)*60*60, 
               labels = scales::label_time(format = "%H:%M")
               ) +
  scale_y_continuous(transform = "symlog", breaks = c(0, 10^(0:5)),
                     labels = c("0", "1", "10", "100", "1 000", "10 000", "100 000")) +
  scale_color_jco() +
  guides(color = "none", linetype = guide_legend("Model")) +
  labs(y = "Melanopic Illuminance (lx, mel EDI)",
       x = NULL) + aes(group = interaction(type, model), linetype = model, col = type) + 
  facet_zoom(xlim = c(5.5,10)*3600) +
  scale_linetype_manual(values = 1:4)

Model_zoom2 <- 
model_data_long %>%
  ggplot(aes(x = time, y = value, group = type, col = type)) +
  geom_point(size = 1.5) +
  geom_line(aes(y = fit)) +
  facet_wrap(~model, ncol = 4) + 
  # geom_line(col = "red") +
  # geom_line(aes(y = fit_m2), col = "skyblue3") +
  # geom_line(aes(y = fit_m3), col = "limegreen") +
  # geom_line(aes(y = fit_m4), col = "orange2") +
  theme_cowplot() +
  scale_x_time(limits = c(0, 24*60*60), expand = c(0,0),
               breaks = c(0, 3, 6, 9, 12, 15, 18, 21, 24)*60*60, 
               labels = scales::label_time(format = "%H:%M")
               ) +
  scale_y_continuous(transform = "symlog", breaks = c(0, 10^(0:5)),
                     labels = c("0", "1", "10", "100", "1 000", "10 000", "100 000")) +
  scale_color_jco() +
  guides(color = "none", linetype = guide_legend("Model")) +
  labs(y = "Melanopic Illuminance (lx, mel EDI)",
       x = NULL) + aes(group = interaction(type, model), linetype = model, col = type) + 
  facet_zoom(xlim = c(18,23.5)*3600) +
  scale_linetype_manual(values = 1:4)

Model_zoom1 + guide_area() + Model_zoom2 + 
  plot_layout(guides = "collect", axes = "collect", widths = c(1, 0.15, 1)) &
  theme(plot.margin = margin(5,10,5,5),
                axis.text = element_text(size = 13),
        axis.title = element_text(size = 16),
        legend.text = element_text(size = 13)
        )
```

```
ggsave("figures/Figure_4.pdf", width = 17.5, height = 8, units = "cm", scale = 2)
```

#### 5 Session info

```
sessionInfo()
```

```
R version 4.4.2 (2024-10-31)
Platform: aarch64-apple-darwin20
Running under: macOS Sequoia 15.1.1

Matrix products: default
BLAS:   /Library/Frameworks/R.framework/Versions/4.4-arm64/Resources/lib/libRblas.0.dylib 
LAPACK: /Library/Frameworks/R.framework/Versions/4.4-arm64/Resources/lib/libRlapack.dylib;  LAPACK version 3.12.0

locale:
[1] en_US.UTF-8/en_US.UTF-8/en_US.UTF-8/C/en_US.UTF-8/en_US.UTF-8

time zone: Europe/Berlin
tzcode source: internal

attached base packages:
[1] stats     graphics  grDevices datasets  utils     methods   base     

other attached packages:
 [1] ggforce_0.4.2     cowplot_1.1.3     ggsci_3.2.0       itsadug_2.4.1    
 [5] plotfunctions_1.4 gt_0.11.1         patchwork_1.3.0   mgcv_1.9-1       
 [9] nlme_3.1-166      lubridate_1.9.4   forcats_1.0.0     stringr_1.5.1    
[13] dplyr_1.1.4       purrr_1.0.2       readr_2.1.5       tidyr_1.3.1      
[17] tibble_3.2.1      ggplot2_3.5.1     tidyverse_2.0.0   LightLogR_0.4.2  

loaded via a namespace (and not attached):
 [1] gtable_0.3.6      xfun_0.49         htmlwidgets_1.6.4 lattice_0.22-6   
 [5] tzdb_0.4.0        vctrs_0.6.5       tools_4.4.2       generics_0.1.3   
 [9] parallel_4.4.2    fansi_1.0.6       pkgconfig_2.0.3   Matrix_1.7-1     
[13] lifecycle_1.0.4   compiler_4.4.2    farver_2.1.2      textshaping_0.4.1
[17] munsell_0.5.1     janitor_2.2.0     snakecase_0.11.1  htmltools_0.5.8.1
[21] yaml_2.3.10       pillar_1.9.0      crayon_1.5.3      MASS_7.3-61      
[25] tidyselect_1.2.1  digest_0.6.37     stringi_1.8.4     labeling_0.4.3   
[29] splines_4.4.2     polyclip_1.10-7   fastmap_1.2.0     grid_4.4.2       
[33] colorspace_2.1-1  cli_3.6.3         magrittr_2.0.3    utf8_1.2.4       
[37] withr_3.0.2       scales_1.3.0      bit64_4.5.2       timechange_0.3.0 
[41] rmarkdown_2.29    ggtext_0.1.2      bit_4.5.0.1       ragg_1.3.3       
[45] hms_1.1.3         evaluate_1.0.1    knitr_1.49        rlang_1.1.4      
[49] gridtext_0.1.5    Rcpp_1.0.13-1     glue_1.8.0        tweenr_2.0.3     
[53] xml2_1.3.6        renv_1.0.11       rstudioapi_0.17.1 vroom_1.6.5      
[57] jsonlite_1.8.9    R6_2.5.1          systemfonts_1.1.0
```


###### Source Code

```
---
title: "Supplement S1"
author: Johannes Zauner
format: 
  html:
    embed-resources: true
    code-tools: true
    toc: true
    toc-depth: 4
    number-sections: true
---

## Preface

this document contains the analysis documentation for the article **How to deal with darkness: Modelling and visualization of zero-inflated personal light exposure data on a logarithmic scale**.

```{r Setup}

### install.packages("devtools")
### devtools::install_github("tscnlab/LightLogR")

library(LightLogR)
library(tidyverse)
library(mgcv)
library(patchwork)
library(gt)
library(itsadug)
library(ggsci)
library(cowplot)
library(ggforce)
```

## Baseline Data and visualization

In this section, we will copy relevant portions of a tutorial from LightLogR [The whole game](https://tscnlab.github.io/LightLogR/articles/Day.html) which will be the basis for both visualizations and analysis.

```{r}
path <- system.file("extdata", 
              package = "LightLogR")

file.LL <- "205_actlumus_Log_1020_20230904101707532.txt.zip"
file.env <- "cyepiamb_CW35_Log_1431_20230904081953614.txt.zip"
tz <- "Europe/Berlin"
dataset.LL <- import$ActLumus(file.LL, path, auto.id = "^(\\d{3})", tz = tz)
dataset.env <- import$ActLumus(file.env, path, manual.id = "CW35", tz = tz)
dataset.LL <- 
  dataset.LL %>% data2reference(Reference.data = dataset.env, across.id = TRUE)
dataset.LL <- 
  dataset.LL %>% select(Id, Datetime, MEDI, Reference)
dataset.LL.partial <- 
dataset.LL %>% #dataset
  filter_Date(start = "2023-09-01", length = days(1)) %>% 
  aggregate_Datetime(unit = "5 mins")
```

## Visualizations

### Pattern

#### Setup

```{r}
day_ribbon <- 
  geom_ribbon(aes(ymin = MEDI, ymax=Reference), 
              alpha = 0.25, fill = "#0073C2FF",
              outline.type = "upper", col = "#0073C2FF", size = 0.15)

participant_ribbon <- 
  geom_ribbon(aes(ymin = 0, ymax = MEDI), alpha = 0.30, fill = "#EFC000", 
              outline.type = "upper", col = "#EFC000", size = 0.4)

lower_bound <- 
 geom_hline(yintercept = 1, col = "red", lty = 3)
upper_bound <- 
 geom_hline(yintercept = 10^5, col = "red", lty = 3)
time_scale <- 
  scale_x_time(breaks = c(0, 6, 12, 18, 24)*3600, labels = 
               scales::label_time(format = "%H:%M"), 
      expand = c(0, 0), limits = c(0, 24 * 3600))

```


#### Visualization A

```{r vis A}

Plot_A <- 
dataset.LL.partial %>% 
   gg_day(facetting = FALSE, geom = "blank", y.scale = "identity",
          x.axis.label = "Local Time (HH:MM)",
          y.axis.breaks = seq(0, to = 10^5, by = 10^4), 
          y.axis.label = "Melanopic Illuminance (lx, mel EDI)") + #base plot
  day_ribbon + participant_ribbon +
 lower_bound + upper_bound + time_scale

Plot_A
```


#### Visualization B

```{r vis B}

Plot_B <- 
dataset.LL.partial %>% 
   gg_day(facetting = FALSE, geom = "blank",
                    x.axis.label = "Local Time (HH:MM)",
          y.axis.label = "Melanopic Illuminance (lx, mel EDI)"
          ) + #base plot
  scale_y_log10(breaks = 10^(-5:5), labels = \(x) paste0("10^",log10(x))) +
  day_ribbon + participant_ribbon +
 lower_bound + upper_bound + time_scale

Plot_B
```

#### Visualization C

```{r vis C}

Plot_C <-
dataset.LL.partial %>% 
  mutate(MEDI = case_when(MEDI == 0 ~ NA,
                          .default = MEDI),
         Reference = case_when(Reference == 0 ~ NA,
                               .default = Reference)) %>% 
   gg_day(facetting = FALSE, geom = "blank", 
                    x.axis.label = "Local Time (HH:MM)",
          y.axis.label = "Melanopic Illuminance (lx, mel EDI)"
          ) + #base plot
    geom_ribbon(
      aes(ymin = 
            ifelse(is.na(MEDI) | MEDI < min(Reference, na.rm = TRUE), 
                   min(Reference, na.rm = TRUE), MEDI), 
          ymax=Reference), 
              alpha = 0.25, fill = "#0073C2FF",
              outline.type = "upper", col = "#0073C2FF", size = 0.15) + #solar reference
  geom_ribbon(aes(ymin = min(MEDI, na.rm = TRUE), ymax = MEDI), alpha = 0.30, 
              fill = "#EFC000",
              outline.type = "upper", col = "#EFC000", size = 0.4) +
  geom_point(
    data = \(x) x %>% filter(is.na(Reference)) %>% mutate(Reference = 10^-4), 
    aes(y = Reference), col = "#0073C2FF", size = 1, alpha = 0.25) +
  geom_point(data = \(x) x %>% filter(is.na(MEDI)) %>% mutate(MEDI = 10^-4), 
              col = "#EFC000", size = 1, alpha = 0.25, position = position_nudge(y=0.05)) +
  scale_y_log10(breaks = 10^(-5:5), 
                labels = \(x) ifelse(x == 10^-4, "0", paste0("10^",log10(x)))
                ) +
 lower_bound + upper_bound + time_scale

Plot_C
```

#### Visualization D

```{r vis D}

Plot_D <-
dataset.LL.partial %>% 
   gg_day(facetting = FALSE, geom = "blank", 
                    x.axis.label = "Local Time (HH:MM)",
          y.axis.label = "Melanopic Illuminance (lx, mel EDI)") + #base plot
 day_ribbon + participant_ribbon +
 lower_bound + upper_bound + time_scale 

Plot_D
```

#### Combined Visualization

```{r}
Pattern <-
Plot_A + plot_spacer() + Plot_B + plot_spacer() + Plot_C + plot_spacer() + Plot_D + 
  plot_layout(ncol = 7, nrow = 1, axes = "collect",
              widths = c(1,-0.15,1,-0.15,1,-0.15,1)) + 
  plot_annotation(tag_levels = "A") & 
  theme(
    plot.tag.position = c(0, 1),
        plot.tag.location = "plot",
        plot.tag = element_text(size = 15, hjust = 0.5, vjust =-0.75),
        axis.text = element_text(size = 13),
        axis.title = element_text(size = 16)
        ) +
  theme(plot.margin = margin(15,10,5,5))

ggsave("figures/Figure_1.pdf", Pattern, width = 17.5, height = 5.5, units = "cm", scale = 2)
```

### Differences

#### Visualization E

```{r vis E}
Plot_E <-
dataset.LL.partial %>% 
   gg_day(y.axis = Reference - MEDI, facetting = FALSE, geom = "blank", 
                    x.axis.label = "Local Time (HH:MM)", y.scale = "identity",
          y.axis.breaks = seq(-10^4, to = 10^5, by = 10^4),
          y.axis.label = "Daylight - Participant (lx, mel EDI)") + #base plot
 geom_area(
   aes(group = consecutive_id((Reference - MEDI) >=0 ),
       fill = (Reference - MEDI) >=0,
       col = (Reference - MEDI) >=0
       ), outline.type = "upper", alpha = 0.25, size = 0.25
 )+
  guides(fill = "none", col = "none") +
  time_scale +
  geom_hline(aes(yintercept = 0), lty = 2, col = "red") + 
    scale_fill_manual(values = c("#EFC000", "#0073C2")) + 
    scale_color_manual(values = c("#EFC000", "#0073C2"))

Plot_E

```

#### Visualization F

```{r vis F}
Plot_F <-
dataset.LL.partial %>% 
   gg_day(facetting = FALSE, geom = "blank", 
                    x.axis.label = "Local Time (HH:MM)",
          y.axis.label = "Daylight - Participant (lx, mel EDI)") + #base plot
 geom_ribbon(
   aes(
     ymin = 0, ymax = abs(Reference - MEDI),
     group = consecutive_id((Reference - MEDI) >=0 ),
       fill = (Reference - MEDI) >=0,
       col = (Reference - MEDI) >=0
       ), outline.type = "upper", alpha = 0.25, size = 0.25
 )+
  guides(fill = "none", col = "none") +
  time_scale +
  geom_hline(aes(yintercept = 0), lty = 2, col = "red") + 
    scale_fill_manual(values = c("#EFC000", "#0073C2")) + 
    scale_color_manual(values = c("#EFC000", "#0073C2")) +
    scale_y_log10(breaks = 10^(-5:5), labels = \(x) paste0("10^",log10(x)))

Plot_F

```

#### Visualization G

```{r vis G}
Plot_G <-
dataset.LL.partial %>% 
  mutate(MEDI = ifelse(Reference-MEDI == 0, NA, MEDI)) %>% 
   gg_day(facetting = FALSE, geom = "blank", 
                    x.axis.label = "Local Time (HH:MM)",
          y.axis.label = "Daylight - Participant (lx, mel EDI)") + #base plot
 geom_ribbon(
   aes(
     ymin = min(abs(Reference-MEDI), na.rm = TRUE), ymax = abs(Reference - MEDI),
     group = consecutive_id((Reference - MEDI) >=0 ),
       fill = (Reference - MEDI) >=0,
       col = (Reference - MEDI) >=0
       ), outline.type = "upper", alpha = 0.25, size = 0.25
 )+
  guides(fill = "none", col = "none") +
  time_scale +
  geom_hline(aes(yintercept = 10^-4), lty = 2, col = "red") + 
    scale_fill_manual(values = c("#EFC000", "#0073C2")) + 
    scale_color_manual(values = c("#EFC000", "#0073C2")) +
  scale_y_log10(breaks = 10^(-5:5), 
                labels = \(x) ifelse(x == 10^-4, "0", paste0("10^",log10(x)))
                ) +
  geom_point(
    data = \(x) x %>% filter(is.na(MEDI)) %>% mutate(MEDI = 10^-4), 
    aes(y = MEDI), col = "grey", size = 1, alpha = 0.5)

Plot_G

```

#### Visualization H

```{r vis H}
Plot_H <-
dataset.LL.partial %>% 
   gg_day(y.axis = Reference - MEDI, facetting = FALSE, geom = "blank", 
                    x.axis.label = "Local Time (HH:MM)",
          y.axis.label = "Daylight - Participant (lx, mel EDI)") + #base plot
 geom_area(
   aes(group = consecutive_id((Reference - MEDI) >=0 ),
       fill = (Reference - MEDI) >=0,
       col = (Reference - MEDI) >=0
       ), outline.type = "upper", alpha = 0.25, size = 0.25
 )+
  guides(fill = "none", col = "none") +
  time_scale +
  geom_hline(aes(yintercept = 0), lty = 2, col = "red") + 
    scale_fill_manual(values = c("#EFC000", "#0073C2")) + 
    scale_color_manual(values = c("#EFC000", "#0073C2"))

Plot_H

```

#### Combined Visualization

```{r}
Differences <- 
Plot_E + plot_spacer() + Plot_F + plot_spacer() + Plot_G + plot_spacer() + Plot_H + 
  plot_layout(ncol = 7, nrow = 1, axes = "collect",
              widths = c(1,-0.15,1,-0.15,1,-0.15,1)) + 
  plot_annotation(tag_levels = "A") & 
  theme(
    plot.tag.position = c(0, 1),
        plot.tag.location = "plot",
        plot.tag = element_text(size = 15, hjust = 0, vjust =-0.75),
        axis.text = element_text(size = 13),
        axis.title = element_text(size = 16)
        ) +
  theme(plot.margin = margin(15,10,5,5))

ggsave("figures/Figure_2.pdf", Differences, width = 17.5, height = 5.5, units = "cm", scale = 2)
```

## Modelling

```{r}
#setting the ends for the cyclic smooth
knots_day <- list(time = c(0, 24*3600))

model_data <- 
dataset.LL.partial %>% 
  ungroup() %>% 
  rename(Environment = Reference) %>% 
  pivot_longer(names_to = "type", cols = c(MEDI, Environment)) %>%  
  arrange(type) %>% 
  mutate(time = hms::as_hms(Datetime) %>% as.numeric(),
         time = c(time[-n()], 24*3600),
         type = factor(type),
         start.event = time == 0,
         input_m1 = 
           case_when(
             value == 0 ~ NA,
             .default = value),
         input_m2 = value + .Machine$double.eps,
         input_m3 = value + 0.1,
         input_m4 = value,
         .by = type
  )
```


### Model type 1: 0 to NA

```{r}
pattern_formula <- log10(input_m1) ~ type + s(time, by = type, bs = "cc", k = 24)

model_data_m1 <- 
  model_data %>% 
  group_by(type, case = is.na(input_m1)) %>% 
  mutate(start.event = 
                        c(TRUE, rep(FALSE, n() -1)))

model_1 <- bam(formula = pattern_formula, 
             data = model_data_m1, 
             method = "REML", 
             knots = knots_day)

r1 <- start_value_rho(model_1, plot=TRUE)

model_1 <- bam(formula = pattern_formula, 
             data = model_data_m1, 
             method = "REML", 
             knots = knots_day, 
             rho = r1,
             AR.start = model_data_m1$start.event
              )

acf_resid(model_1)

summary(model_1)

gam.check(model_1, rep = 500)

estimate_m1 <- 
plot_smooth(model_1, 
            view = "time", 
            plot_all = "type",
            # rug = F,
            n.grid = 12*24+1,
            rm.ranef = FALSE,
            se = 0)

estimate_m1_dif <- 
plot_diff(model_1,
          view = "time",
          comp = list(type = c("Environment", "MEDI")),
          rm.ranef = FALSE,
          n.grid = 12*24+1)

estimate_m1$fv %>% 
  ggplot(aes(x = time, y = 10^fit, col = type)) +
  geom_line() +
  theme_minimal() +
  scale_x_time(limits = c(0, 24*60*60), expand = c(0,0),
               breaks = c(0, 6, 12, 18, 24)*60*60, 
               labels = scales::label_time(format = "%H:%M")
               ) +
  scale_y_continuous(transform = "symlog", breaks = c(0, 10^(0:5))) +
  geom_point(data = model_data, aes(y = value, col = type))

model_data <- 
model_data %>% 
  left_join(estimate_m1$fv, by = c("type", "time")) %>% 
  rename(fit_m1 = fit) %>% 
  mutate(fit_m1 = (10^(fit_m1)))

```

### Model type 2: adding smallest float to every value

```{r}
pattern_formula <- log10(input_m2) ~ type + s(time, by = type, bs = "cc", k = 24)

model_2 <- bam(formula = pattern_formula, 
             data = model_data, 
             method = "REML", 
             knots = knots_day)

r1 <- start_value_rho(model_2, plot=TRUE)

model_2 <- bam(formula = pattern_formula, 
             data = model_data, 
             method = "REML", 
             knots = knots_day, 
             rho = r1,
             AR.start = model_data$start.event
              )

acf_resid(model_2)

summary(model_2)

gam.check(model_2, rep = 500)

estimate_m2 <- 
plot_smooth(model_2, 
            view = "time", 
            plot_all = "type",
            # rug = F,
            n.grid = 12*24+1,
            rm.ranef = FALSE,
            se = 0)

estimate_m2_dif <- 
plot_diff(model_2,
          view = "time",
          comp = list(type = c("Environment", "MEDI")),
          rm.ranef = FALSE,
          n.grid = 12*24+1)

estimate_m2$fv %>% 
  ggplot(aes(x = time, y = (10^fit)-.Machine$double.eps, col = type)) +
  geom_line() +
  theme_minimal() +
  scale_x_time(limits = c(0, 24*60*60), expand = c(0,0),
               breaks = c(0, 6, 12, 18, 24)*60*60, 
               labels = scales::label_time(format = "%H:%M")
               ) +
  scale_y_continuous(transform = "symlog", breaks = c(0, 10^(0:5))) +
  geom_point(data = model_data, aes(y = value, col = type))

model_data <- 
model_data %>% 
  left_join(estimate_m2$fv, by = c("type", "time")) %>% 
  rename(fit_m2 = fit) %>% 
  mutate(fit_m2 = (10^(fit_m2)) - .Machine$double.eps)

```
### Model type 3: adding -1 log10 to every value

```{r}
pattern_formula <- log10(input_m3) ~ type + s(time, by = type, bs = "cc", k = 24)

model_3 <- bam(formula = pattern_formula, 
             data = model_data, 
             method = "REML", 
             knots = knots_day)

r1 <- start_value_rho(model_3, plot=TRUE)

model_3 <- bam(formula = pattern_formula,
             data = model_data,
             method = "REML",
             knots = knots_day,
             rho = r1,
             AR.start = model_data$start.event
              )

acf_resid(model_3)

summary(model_3)

gam.check(model_3, rep = 500)

estimate_m3 <- 
plot_smooth(model_3, 
            view = "time", 
            plot_all = "type",
            # rug = F,
            n.grid = 12*24+1,
            rm.ranef = FALSE,
            se = 0)

estimate_m3_dif <- 
plot_diff(model_3,
          view = "time",
          comp = list(type = c("Environment", "MEDI")),
          rm.ranef = FALSE,
          n.grid = 12*24+1)

estimate_m3$fv %>% 
  ggplot(aes(x = time, y = (10^fit)-0.1, col = type)) +
  geom_line() +
  theme_minimal() +
  scale_x_time(limits = c(0, 24*60*60), expand = c(0,0),
               breaks = c(0, 6, 12, 18, 24)*60*60, 
               labels = scales::label_time(format = "%H:%M")
               ) +
  scale_y_continuous(transform = "symlog", breaks = c(0, 10^(0:5))) +
  geom_point(data = model_data, aes(y = value, col = type))

model_data <- 
model_data %>% 
  left_join(estimate_m3$fv, by = c("type", "time")) %>% 
  rename(fit_m3 = fit) %>% 
  mutate(fit_m3 = (10^(fit_m3))-0.1)

```

### Model type 4: using the Tweedy family

```{r}
pattern_formula <- input_m4 ~ type + s(time, by = type, bs = "cc", k = 24)

model_4 <- gam(formula = pattern_formula, 
             data = model_data, 
             method = "REML", 
             family = tw,
             knots = knots_day)

summary(model_4)

gam.check(model_4, rep = 500)

plot(model_4)

estimate_m4 <- 
plot_smooth(model_4, 
            view = "time", 
            plot_all = "type",
            # rug = F,
            n.grid = 12*24+1,
            rm.ranef = FALSE,
            se = 0)

estimate_m4_dif <- 
plot_diff(model_4,
          view = "time",
          comp = list(type = c("Environment", "MEDI")),
          rm.ranef = FALSE,
          n.grid = 12*24+1)

estimate_m4$fv %>% 
  ggplot(aes(x = time, y = exp(fit), col = type)) +
  geom_line() +
  theme_minimal() +
  scale_x_time(limits = c(0, 24*60*60), expand = c(0,0),
               breaks = c(0, 6, 12, 18, 24)*60*60, 
               labels = scales::label_time(format = "%H:%M")
               ) +
  scale_y_continuous(transform = "symlog", breaks = c(0, 10^(0:5))) +
  geom_point(data = model_data, aes(y = value, col = type))

model_data <- 
model_data %>% 
  left_join(estimate_m4$fv, by = c("type", "time")) %>% 
  rename(fit_m4 = fit) %>% 
  mutate(fit_m4 = exp(fit_m4))

```

### Combined Models

```{r combining models}
#adding scaled.pearson residuals
model_data$resid_m1 <- NA
model_data$resid_m1[!is.na(model_data$input_m1)] <- 
  (residuals(model_1, type = "scaled.pearson"))
model_data$resid_m2 <-(residuals(model_2, type = "scaled.pearson"))
model_data$resid_m3 <- (residuals(model_3, type = "scaled.pearson"))
model_data$resid_m4 <- (residuals(model_4, type = "scaled.pearson"))

model_data_long <- 
model_data %>% 
  pivot_longer(cols = input_m1:resid_m4, names_to = c(".value", "model"), 
               names_pattern = "(.*)_(m\\d{1})") %>% 
  mutate(model = model %>% str_replace(pattern = "m", replacement =  "Model "))


summary(model_1)
summary(model_2)
summary(model_3)
summary(model_4)
```


```{r combining models2}
#model fitted plot
Model_fitted_plot <- 
model_data_long %>%
  ggplot(aes(x = time, y = value, group = type, col = type)) +
  geom_point(size = 0.5) +
  geom_line(aes(y = fit), col = "black") +
  facet_wrap(~model, ncol = 4) + 
  # geom_line(col = "red") +
  # geom_line(aes(y = fit_m2), col = "skyblue3") +
  # geom_line(aes(y = fit_m3), col = "limegreen") +
  # geom_line(aes(y = fit_m4), col = "orange2") +
  theme_cowplot() +
  scale_x_time(limits = c(0, 24*60*60), expand = c(0,0),
               breaks = c(0, 6, 12, 18, 24)*60*60, 
               labels = scales::label_time(format = "%H:%M")
               ) +
  scale_y_continuous(transform = "symlog", breaks = c(0, 10^(0:5)),
                     labels = c("0", "1", "10", "100", "1 000", "10 000", "100 000")) +
  scale_color_jco() +
  guides(color = "none") +
  labs(y = "Melanopic Illuminance\n(lx, mel EDI)",
       x = NULL) +
  theme(
        strip.text = element_text(size = 14)
  )
 

Model_residual_plot <- 
model_data_long %>%   
ggplot(aes(x=time))+
  geom_point(aes(y=resid), size = 0.5) +
  facet_wrap(~model, ncol = 4) +
  theme_cowplot() +
  coord_cartesian(ylim = c(-5,5)) +
  time_scale +
  labs(y = "Scaled Pearson\nresiduals", x = "Local time (HH:MM)") +
  theme(strip.background = element_blank(),
        strip.text = element_blank())

Model_fitted_plot / Model_residual_plot + 
  plot_layout(guides = "collect", heights = c(3,2)) +
  plot_annotation(tag_levels = "A") &
  theme(
    plot.tag.position = c(0, 0.99),
    plot.tag = element_text(size = 15, hjust = 0.5, vjust = 0, face = "bold"),
    panel.spacing = unit(2.5, "lines"),
        plot.tag.location = "plot",
        axis.text = element_text(size = 13),
        axis.title = element_text(size = 16)
        ) +
  theme(plot.margin = margin(15,10,5,5))

  ggsave("figures/Figure_3.pdf", width = 17.5, height = 8, units = "cm", scale = 2)
  
```


```{r combining models3}

Model_zoom1 <- 
model_data_long %>%
  ggplot(aes(x = time, y = value, group = type, col = type)) +
  geom_point(size = 1.5) +
  geom_line(aes(y = fit)) +
  facet_wrap(~model, ncol = 4) + 
  # geom_line(col = "red") +
  # geom_line(aes(y = fit_m2), col = "skyblue3") +
  # geom_line(aes(y = fit_m3), col = "limegreen") +
  # geom_line(aes(y = fit_m4), col = "orange2") +
  theme_cowplot() +
  scale_x_time(limits = c(0, 24*60*60), expand = c(0,0),
               breaks = c(0, 3, 6, 9, 12, 15, 18, 21, 24)*60*60, 
               labels = scales::label_time(format = "%H:%M")
               ) +
  scale_y_continuous(transform = "symlog", breaks = c(0, 10^(0:5)),
                     labels = c("0", "1", "10", "100", "1 000", "10 000", "100 000")) +
  scale_color_jco() +
  guides(color = "none", linetype = guide_legend("Model")) +
  labs(y = "Melanopic Illuminance (lx, mel EDI)",
       x = NULL) + aes(group = interaction(type, model), linetype = model, col = type) + 
  facet_zoom(xlim = c(5.5,10)*3600) +
  scale_linetype_manual(values = 1:4)

Model_zoom2 <- 
model_data_long %>%
  ggplot(aes(x = time, y = value, group = type, col = type)) +
  geom_point(size = 1.5) +
  geom_line(aes(y = fit)) +
  facet_wrap(~model, ncol = 4) + 
  # geom_line(col = "red") +
  # geom_line(aes(y = fit_m2), col = "skyblue3") +
  # geom_line(aes(y = fit_m3), col = "limegreen") +
  # geom_line(aes(y = fit_m4), col = "orange2") +
  theme_cowplot() +
  scale_x_time(limits = c(0, 24*60*60), expand = c(0,0),
               breaks = c(0, 3, 6, 9, 12, 15, 18, 21, 24)*60*60, 
               labels = scales::label_time(format = "%H:%M")
               ) +
  scale_y_continuous(transform = "symlog", breaks = c(0, 10^(0:5)),
                     labels = c("0", "1", "10", "100", "1 000", "10 000", "100 000")) +
  scale_color_jco() +
  guides(color = "none", linetype = guide_legend("Model")) +
  labs(y = "Melanopic Illuminance (lx, mel EDI)",
       x = NULL) + aes(group = interaction(type, model), linetype = model, col = type) + 
  facet_zoom(xlim = c(18,23.5)*3600) +
  scale_linetype_manual(values = 1:4)

Model_zoom1 + guide_area() + Model_zoom2 + 
  plot_layout(guides = "collect", axes = "collect", widths = c(1, 0.15, 1)) &
  theme(plot.margin = margin(5,10,5,5),
                axis.text = element_text(size = 13),
        axis.title = element_text(size = 16),
        legend.text = element_text(size = 13)
        )

ggsave("figures/Figure_4.pdf", width = 17.5, height = 8, units = "cm", scale = 2)

```

## Session info

```{r}
sessionInfo()
```
```
